# Bayesian Inference of Pathogen Phylogeography using the Structured Coalescent Model

**DOI:** 10.1101/2024.10.14.617553

**Authors:** Ian Roberts, Richard G. Everitt, Jere Koskela, Xavier Didelot

## Abstract

Over the past decade, pathogen genome sequencing has become well established as a powerful approach to study infectious disease epidemiology. In particular, when multiple genomes are available from several geographical locations, comparing them is informative about the relative size of the local pathogen populations as well as past migration rates and events between locations. The structured coalescent model has a long history of being used as the underlying process for such phylogeographic analysis. However, the computational cost of using this model does not scale well to the large number of genomes frequently analysed in pathogen genomic epidemiology studies. Several approximations of the structured coalescent model have been proposed, but their effects are difficult to predict. Here we show how the exact structured coalescent model can be used to analyse a precomputed dated phylogeny, in order to perform Bayesian inference on the past migration history, the effective population sizes in each location, and the directed migration rates from any location to another. We describe an efficient reversible jump Markov Chain Monte Carlo scheme which is implemented in a new R package. We use simulations to demonstrate the scalability and correctness of our method and to compare it with existing comparable software. We also applied our new method to several state-of-the-art datasets on the population structure of real pathogens to showcase the relevance of our method to current data scales and research questions.

## INTRODUCTION

Genomic data is readily available for a wide range of pathogens from large online databases (Benson et al., 2017; Shu and McCauley, 2017; Jolley et al., 2018), and can be obtained relatively cheaply and quickly for newly acquired clinical samples (Didelot et al., 2012; Houldcroft et al., 2017; Gardy and Loman, 2018). Comparative analysis of these genomic sequences can help understand the way pathogens cause disease, spread and evolve, and this general approach is called pathogen phylodynamics (Grenfell et al., 2004; Volz et al., 2013; Baele et al., 2016). A specific problem in phylodynamics is to try and understand the population geographical structure given genomes from several distinct locations. This approach is called pathogen phylogeography, and typically attempts to infer the relative size of the pathogen populations in each location, as well as the rates of migrations between them (Pybus and Rambaut, 2009; Bloomquist et al., 2010; Grubaugh et al., 2019).

Under many forward-in-time population genetics models, including the Wright–Fisher model (Fisher, 1922; Wright, 1931), the Moran model (Moran, 1958) and more generally the Cannings models family (Cannings, 1974), the ancestry of a sample from an unstructured population is described by the coalescent model (Kingman, 1982a,b). On the other hand if the population is structured following a finite island model with migrations (Wright, 1931), then the ancestry of samples from several locations is described by an extension of the coalescent model called the structured coalescent model. This model was first presented in the case with two locations (Takahata, 1988), then extended to the general case (Notohara, 1990) and its properties have been subsequently thoroughly examined (Hudson, 1990; Herbots, 1994; Donnelly and Tavaré, 1995; Nordborg, 1997). Here we consider the structured coalescent model on a natural time scale and with leaves sampled over time (Drummond et al., 2002), which enables the study of measurably evolving populations (Drummond et al., 2003; Biek et al., 2015).

Inference under the structured coalescent model is challenging due to the mixed discrete and continuous components required to fully specify a structured genealogy, the high dimensionality of the latent genealogy, multimodality, potential for heavy tails in posteriors, complicated correlation structures, etc. Current inference methods for structured genealogies fall into two broad categories, using Markov chain Monte Carlo (MCMC) to sample full migration histories under the exact structured coalescent model, or integrating out parts of the migration history by using an approximation. Exact inference methods often fail to scale to large datasets, and are computationally demanding. Approximate methods improve the scalability and can be applied to larger datasets. By far the most widely used approach to phylogeography is discrete trait analysis (DTA) in which the geographical location is modelled as evolving independently along the branches of a genealogy in a similar manner to mutation on a genetic locus (Lemey et al., 2009; Gao et al., 2023). It was not originally presented as such, but DTA can be thought of as a rough approximation of the structured coalescent model (De Maio et al., 2015). Less approximate models have been proposed and implemented for example in BASTA (De Maio et al., 2015) and MASCOT (Müller et al., 2018), although at the cost of being much less scalable. These approximate models rely to some extent on ignoring correlation between the current state of contemporary lineages. However, applying these approximate models to data assumed to arise under the structured coalescent can lead to unpredictable biases to the inference (Müller et al., 2017). Furthermore, integrating out parts of the migration history means that these methods cannot directly answer some simple questions of interest, such as how many migration events took place in the ancestry. We therefore focus here on the problem of performing inference under the exact structured coalescent model.

A first MCMC scheme to perform Bayesian inference under the structured coalescent was proposed two decades ago by Ewing et al. (2004), which extended MCMC operators used for inference under the unstructured coalescent (Drummond et al., 2002) as well as introducing new MCMC operators which act directly on migration histories. These migration history operators add or remove migration events from a migration history singly or in pairs. This scheme is irreducible over the space of structured genealogies but suffers from poor performance with slow exploration of the target space. The authors’ implementation of the code is no longer available, however it has been reimplemented at least once before by Vaughan et al. (2014), and we have also reimplemented it in the current study. MultiTypeTree (Vaughan et al., 2014) is a package of BEAST2 (Bouckaert et al., 2019) that also extends the unstructured coalescent operators (Drummond et al., 2002) to the structured coalescent and introduces an additional, more sophisticated node retype operator which modifies the migration history along branches of the genealogy surrounding a coalescent node. Since MultiTypeTree uses the unapproximated structured coalescent model, it has been used as a gold standard against which approximate methods have been compared, but it is computationally demanding and cannot handle large datasets (De Maio et al., 2015; Müller et al., 2017).

In order to ensure that our new phylogenetic approach can scale to the large datasets currently available in pathogen genomics, we separate the inference of a phylogeny and associated migration histories (Didelot and Parkhill, 2022). A dated phylogeny is previously inferred from genomic data, either directly for example using BEAST (Suchard et al., 2018) or BEAST2 (Bouckaert et al., 2019), or by inferring dates from an undated phylogeny using methods such as LSD (To et al., 2016), treedater (Volz and Frost, 2017), TreeTime (Sagulenko et al., 2018) or BactDating (Didelot et al., 2018). Migration histories and the evolutionary parameters governing the structured coalescent model are therefore inferred conditional on this fixed dated phylogeny and the known sampling locations of the genomes. Both previously proposed MCMC schemes (Ewing et al., 2004; Vaughan et al., 2014) contain operators which can be used to sample migration histories for a fixed phylogeny but neither scheme is optimised for this purpose since they both considered the joint problem of inference of the phylogeny and phylogeography. We present a new MCMC scheme optimised for this purpose, which updates the migration history using as proposal distribution a localised, conditional version of DTA (Lemey et al., 2009). We apply our new method to many simulated and real datasets to demonstrate its correctness, scalability and usefulness.

## METHODS

### The Structured Coalescent

The genealogy of a finite sample of homologous, non-recombinant individuals drawn from a larger population is widely modelled using the Kingman coalescent (Kingman, 1982a,b). Under the Kingman coalescent, each pair of contemporary lineages is subject to a coalescent process whereby a pair of lineages coalesce at a fixed rate *θ* = (*N*_e_*g*)^−1^, dependent on the effective population size (*N*_e_) and generation length (*g*) of the population. A natural extension to the Kingman coalescent is to introduce population structure using an island model (Wright, 1931), in which each lineage is assigned to one of finitely-many demes, or subpopulations, at every time. This extension is referred to as the structured coalescent (Hudson, 1990; Donnelly and Tavaré, 1995; Nordborg, 1997).

Under the structured coalescent, lineages are subject to three event types - sampling, coalescence and migration. A sampling event corresponds to adding a new lineage to the process, represented as a new leaf in the phylogenetic tree with a known isolation time and sampling deme. Coalescence events correspond to a pair of contemporary lineages in the same deme finding a common ancestor and coalescing into a single lineage. Coalescence events in deme *i* occur at a fixed rate *θ*_*i*_, which is equivalent to the Kingman coalescent rate for an unstructured population with effective population size equal to that of deme *i*. Migration events correspond to a single lineage moving from its current deme into a new deme. Migration events from deme *i* into deme *j* (*j* ≠ *i*) occur at a fixed rate *λ*_*ij*_ backwards-in-time. A structured genealogy is constructed by simultaneously tracing each of these event types backwards-in-time until the most recent common ancestor (MRCA) of the entire sample is reached. Once complete, the structured genealogy consists of a phylogenetic tree 𝒯 superimposed with a migration history ℋ giving the deme membership at every point on 𝒯.

The structured coalescent probability density can be constructed recursively by considering the time increments between consecutive events (migration, coalescent or sampling). The *r*^th^ time increment of length *τ*_*r*_ contributes a factor

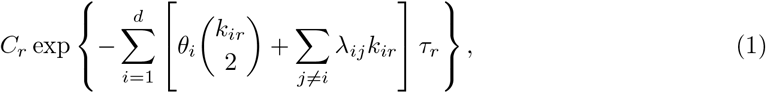

where the constant *C*_*r*_ depends on the leaf-wards event of the interval,

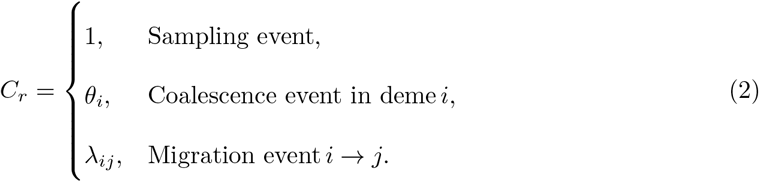

The full probability density for a structured genealogy 𝒢 can then be factorised as

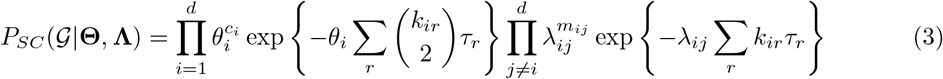

where *c*_*i*_ denotes the total number of coalescence events occurring in deme *i, m*_*ij*_ denotes the total number of migration events from deme *i* to deme *j* backwards-in-time and *k*_*ir*_ denotes the total number of lineages in deme *i* during time increment *r*.

### Discrete Trait Analysis

The Discrete Trait Analysis (DTA) method (Lemey et al., 2009) is a widely-used phylogeographic model which can be viewed as an approximation to the structured coalescent. Under DTA, migration events are placed on a phylogenetic tree as the points of a marked Poisson point process, arising from a forwards-in-time migration process propagating demes leaf-wards through the tree. Whilst this approximation incurs a substantially smaller computational cost than the structured coalescent when performing likelihood-based inference, treating the migration and coalescent processes independently can lead to biased estimates of migration rates (De Maio et al., 2015).

Under the DTA model, a structured genealogy is constructed in two phases by first sampling a dated phylogeny 𝒯 using the Kingman coalescent, and superimposing with a migration history ℋ A migration history is obtained by sampling a deme at the root of the phylogeny and propagating forwards-in-time towards the leaves. The deme propagating through the tree evolves according to a continuous-time Markov process with transition rates given by the forwards-in-time migration rate matrix **F**. When a coalescent event is reached, the migration process splits into two independent copies which continue evolving along each child branch. The DTA probability density of a structured genealogy 𝒢 with *n* samples from *d* demes is given by

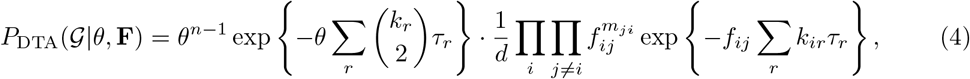

where **F** = (*f*_*ij*_) denotes the matrix of forwards-in-time migration rates, *m*_*ij*_ denotes the total number of migration events from deme *i* to deme *j backwards-in-time* and *k*_*r*_ = Σ _*i*_ *k*_*ir*_ denotes the number of contemporary lineages in the *r*^th^ time increment of length *τ*_*r*_. The forwards-in-time migration rates matrix which maintains the population flow forwards- and backwards-in-time when compared to a backwards-in-time migration process with rates matrix Λ and coalescent rates vector Θ can be computed using (Bahlo and Griffiths, 2000)

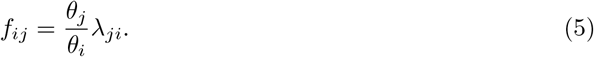

Note that the parameter *θ* in Equation (4) represents the coalescent rate for the entire population considered together, and does not relate to the coalescent rates vector Θ in the structured coalescent.

### Bayesian Inference

Our inferential target is the joint posterior distribution of migration histories ℋ, backwards-in-time migration rates matrices **Λ**and coalescent rates vectors ***θ*** given a fixed phylogenetic tree 𝒯. We assume throughout that the sampling deme of each sampled lineage is known and fixed, with all relevant distributions implicitly conditioned on these fixed demes. The joint posterior distribution can then be specified as

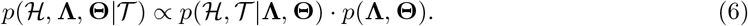

The first term on the right-hand side gives the joint probability of the phylogenetic tree and migration history conditional on the coalescent rates vector **Θ** and migration rates matrix **Λ** and is computed using the structured coalescent probability density (3). The second term gives the prior distribution on the evolutionary parameters.

The specification of prior distributions is a vital part of any Bayesian analysis, and is especially important for inference under the structured coalescent. Several currently-published inference methods, including DTA and MultiTypeTree, use lognormal priors on both migration rates and effective population sizes with parameters *µ* = 0 and *σ* = 4. However, these priors are sometimes criticised for making strong, biologically unreasonable assumptions about the underlying migration and coalescent processes (Gao et al., 2023) and we hence develop our own set of default priors.

We use Gamma-distributed priors for our default framework, with each coalescent rate and backwards-in-time migration rate assigned an independent, and potentially unique, prior distribution. The Gamma distribution describes a flexible family of distributions with options to construct relatively diffuse, uninformative prior distributions as well as more concentrated prior distributions focussing posterior sampling into smaller regions of the target space. Gamma priors are also conjugate with the structured coalescent likelihood and permit exact conditional distributions to be computed and consequently Gibbs samplers can be used to update evolutionary parameters.

Instead of specifying a single default prior distribution, we tailor our default priors to a specific dated phylogeny under four assumptions. We assign independent and identically distributed (i.i.d.) priors to each coalescent rate and a potentially different i.i.d. prior to each backwards-in-time migration rate and reduce the number of prior parameters by setting the shape parameter to *α* = 1, equivalently placing exponentially-distributed priors on evolutionary parameters. We then match the prior expectation with the evolutionary parameters maximising an approximation to the structured coalescent probability density (3) using the following four assumptions to remove dependence on a specific migration history:

i. All demes have equal coalescent rates, *θ*_*i*_ = *θ* for all *i* = 1, 2,…, *d*
ii. All migration rates are equal, *λ*_*ij*_ = *λ* for all *i* ≠ *j*
iii. Lineages are uniformly distributed between demes at every time, 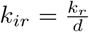 where *k*_*ir*_ denotes the number of lineages in deme *i* during time increment *r* and 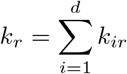 denotes the total number of lineages during time increment *r*
iv. The migration history is parsimonious and contains the minimum number of required migration events 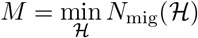

Under assumptions (i)-(iv) the structured coalescent probability density satisfies

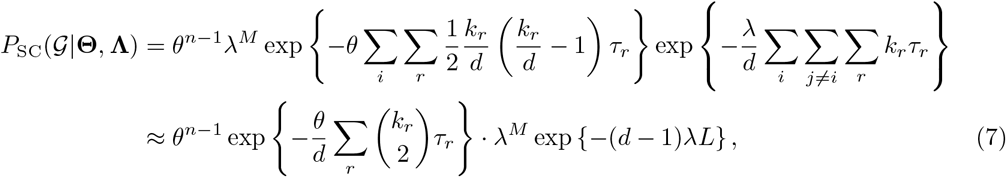

which is maximised by parameter values 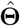 and **Λ**given elementwise by

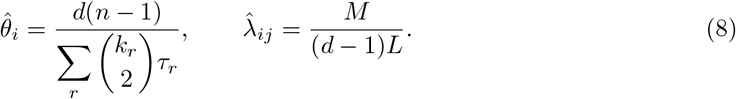

Matching the prior expectations with the maximisers (8) hence yields default prior distributions

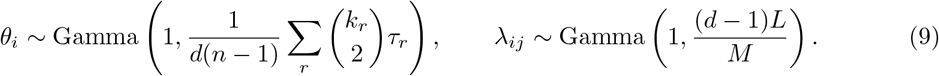

We complete our specification of prior distributions by computing *M*, the minimum number of required migration events for phylogeny 𝒯 using the Fitch algorithm (Fitch, 1971).

### Markov Chain Monte Carlo

We draw joint samples (ℋ, **Θ**, Λ) of migration histories and evolutionary parameters from the posterior distribution (6) using Markov Chain Monte Carlo (MCMC) sampling. An irreducible MCMC scheme is described consisting of three MCMC operators with a Gibbs update for coalescent rate vector **Θ**, a Gibbs update for backwards-in-time migration rate matrix **Λ**, and a Metropolis– Hastings update for migration history ℋ.

#### Evolutionary Parameter Updates

We sample updates to evolutionary parameters using two types of Gibbs update, one for coalescent rates and one for backwards-in-time migration rates. Gamma-distributed prior distributions are conjugate with the structured coalescent for both coalescent rates and backwards-in-time migration rates, hence allowing conditional distributions to be evaluated analytically. We sample updates to evolutionary parameters from their respective Gamma-distributed conditional distributions

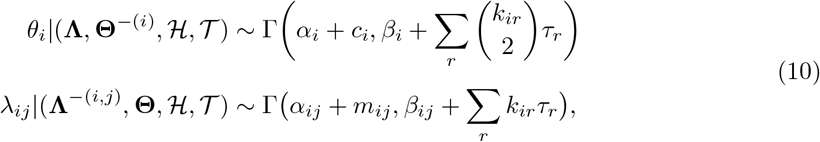

where *c*_*i*_ denotes the number of coalescent events in deme *i* and *m*_*ij*_ denotes the number of backwards-in-time migration events from deme *i* into deme *j* in the structured genealogy 𝒢.

#### Migration History Updates

Joint samples (**Θ, Λ**, ℋ) of evolutionary parameters and migration histories ℋ also require a proposal mechanism to update the migration history conditional on the current evolutionary parameters and fixed dated phylogeny. We describe a family of proposal mechanisms using a localised version of DTA to generate updates of the migration history, which we refer to as Local DTA (Algorithm 1). A Local DTA update samples a subtree of the fixed dated phylogeny on which to update the migration history and then simulates the migration process under DTA conditional on the demes at the points the subtree reconnects to the remainder of the phylogeny. This process relies on the fact that under DTA, migration histories along branches of a phylogeny are conditionally independent given the demes at internal coalescent events and lineage sampling demes. We can hence construct a Local DTA sample sequentially by first sampling the deme at any internal coalescent events of the subtree and subsequently sampling migration histories along branch segment.

Figure 1 illustrates a Local DTA proposal for a phylogeny with three leaves. In the first panel, the subtree highlighted in grey is sampled and the migration history for this subtree is removed. In the second panel, the deme at the MRCA (the only internal coalescent event) is updated conditional on the demes at the points where the subtree reconnects with the remainder of the tree before sampling new migration histories along each of the child branches of the MRCA in the third panel. On the left-hand branch, the subtree reconnects in the red deme between *I* and *R* with no migration events, and on the right-hand branch the subtree reconnects in the blue deme between 3 and *R*, adding a single migration event. The completed proposal has removed two migration events (both from the red deme into the green deme backwards-in-time) and shifted the migration event from the blue deme to the red deme closer to the root of the tree. In the following sections, we highlight the mechanism used to select a subtree, update the deme at the internal coalescent event and sample each of the migration histories.

**Figure 1:**
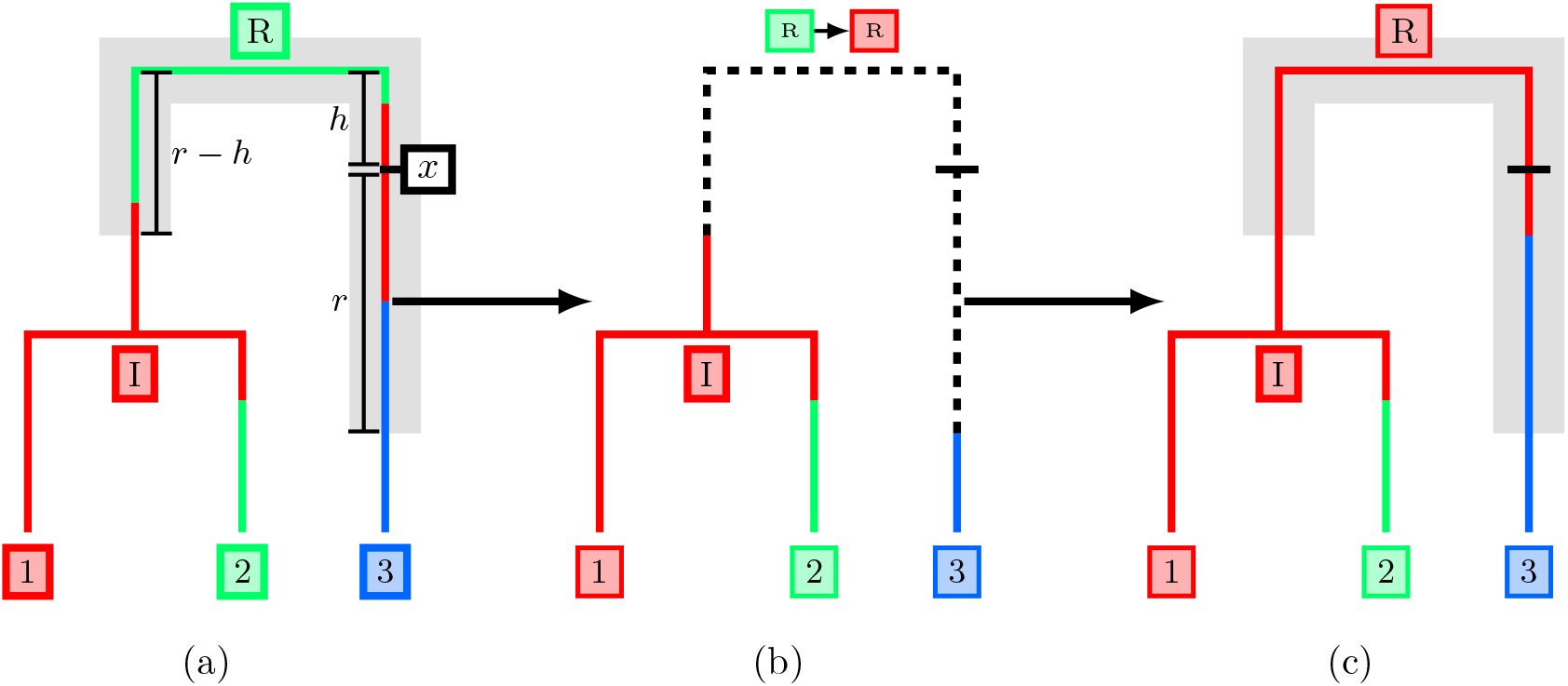
Full migration history update using a radius-based subtree selection. A subtree is sampled centred at the point *x* with radius *r* (a). The migration history on the subtree is erased and the deme at coalescent event *R* is resampled (b). Finally, a new migration history is sampled under a conditional DTA model given the demes at the points where the subtree reconnects to the migration history (c).

##### Algorithm 1

Local DTA

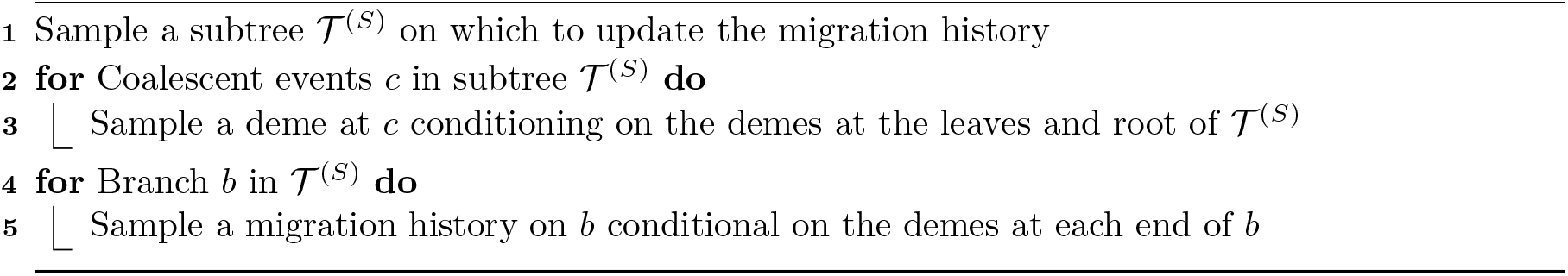

### Subtree Selection

The choice of a subtree selection method directly affects the computational complexity of Local DTA updates. Migration history updates associated with large subtrees containing many coalescent events and a greater total branch length are generally more ambitious, potentially making large changes to the migration history, and are more computationally demanding to construct. We have settled on a radius-based approach in which we include points in the subtree based on their patristic distance from a sampled subtree centre (marked *x* in Figure 1a). We define the patristic distance between two points on a tree to be the sum of branch lengths along the unique path connecting the two points. A subtree of radius *r* is then constructed in two steps, first selecting a location *x* on the underlying tree for the subtree centre and then isolating all points at most patristic distance *r* from *x*. We sample the location of the subtree centre *x* uniformly between the age of the subtree centre between the age of the MRCA, *t*_MRCA_, and the most recently sampled leaf, *t*_Max_, and then select one of the coexisting lineages at that time uniformly at random.

We use a Robbins-Monro Controlled MCMC approach (Robbins and Monro, 1951; Andrieu and Robert, 2001; Andrieu and Thoms, 2008) to adapt the radius *r* during the MCMC run based on the acceptance rate of migration history proposals. The adaptation begins with an initial radius *r*_0_ a target acceptance probability *α*^*^ ∈ (0, 1) and adaptation rate *β* ∈ (1*/*2, 1]. We typically choose an initial radius around one tenth of the total tree height, target acceptance rate *α*^*^ = 0.234 and adaptation rate *β* = 0.6. An acceptance rate of 0.234 is optimal in a range of scenarios, especially when the target distribution follows a Gaussian distribution (Gelman et al., 1996). Whilst our migration history target certainly does not follow a Gaussian distribution and we do not expect this value to be optimal for our MCMC proposal, we find that this value strikes a reasonable balance between exploring the space rapidly and the radius remaining at a reasonable size. The radius *r*_*n*_ at iteration *n* is then updated on a log scale by the update rule

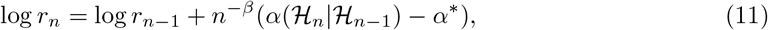

where *α*(ℋ_*n*_|ℋ_*n*−1_) denotes the acceptance probability of a migration history ℋ’ sampled from current state ℋ.

### Coalescent Node Sampling

After selecting a subtree, we update the deme at each coalescent event contained within the subtree, conditioning on the current deme at each point where the subtree reconnects to the remainder of the migration history. These fixed demes correspond to the leaves of the subtree, and the root of the subtree provided the subtree root is not the MRCA. We use belief propagation (Pearl, 1982; Lauritzen and Spiegelhalter, 1988) to compute the conditional distributions at each coalescent node.

In place of computing a full conditional distribution at each coalescent event, we draw samples from the joint distribution of demes using a backward filtering-forward sampling approach similar to the forward-backward algorithm for Hidden Markov Models (Rabiner, 1989). We describe this approach applied to a tree containing only fixed leaf demes, however this can be extended to subtrees where the root deme is also fixed by passing a single message from the subtree root to its children.

The backward filtering pass begins at the leaves of the subtree and passes messages towards the root along all parent edges, following Felsenstein’s pruning algorithm (Felsenstein, 1981). Each message consists of a vector of unnormalised marginal probabilities that the parent node falls into each deme. Formally, a node *i* passes a message *µ*_*ij*_ to its parent *j* consisting of elements

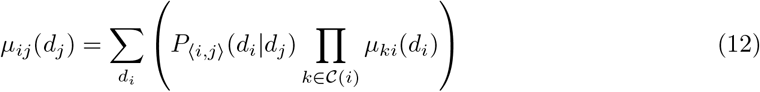

where 𝒞 (*i*) denotes the set of child nodes of *i, d*_*_ denotes the deme at node * and *P*_⟨*i,j*⟩_(*y*|*x*) denotes the transition probability of migrating forward-in-time from deme *x* to deme *y* along edge ⟨*i, j*⟩. A leaf node *i* with a fixed deme has no incoming messages and simply passes the message

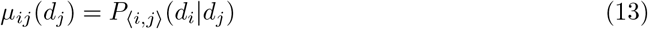

to its parent *j*. All incoming messages to a node must be determined before a message can be passed onwards, which is achieved by computing messages in time order, beginning at the most recent leaf of the subtree and ending at the eldest child of the subtree root. Once all messages have been passed to the root of the tree, the conditional deme distribution at the root is computed by taking the product of all incoming messages.

We begin the forward sampling pass by sampling from the conditional distribution at the root. The root is then treated as fully determined and passes messages to its child nodes using Equation (13). The child nodes now have all incoming messages fully determined and the conditional deme distribution can be computed by taking a product of incoming messages. This process is repeated iteratively until a new deme has been sampled at every coalescent event in the tree. Figure 2 illustrates the messages required for a full pass of the backward filtering-forward sampling algorithm. The demes are computed in the order green-orange-red and the deme distributions at *R* and *I* are sampled from

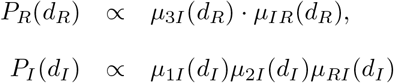

respectively. The messages in grey would be given by a point mass as the transition probability from the determined parent to the determined child along the edge but are unnecessary to sample a deme configuration.

**Figure 2:**
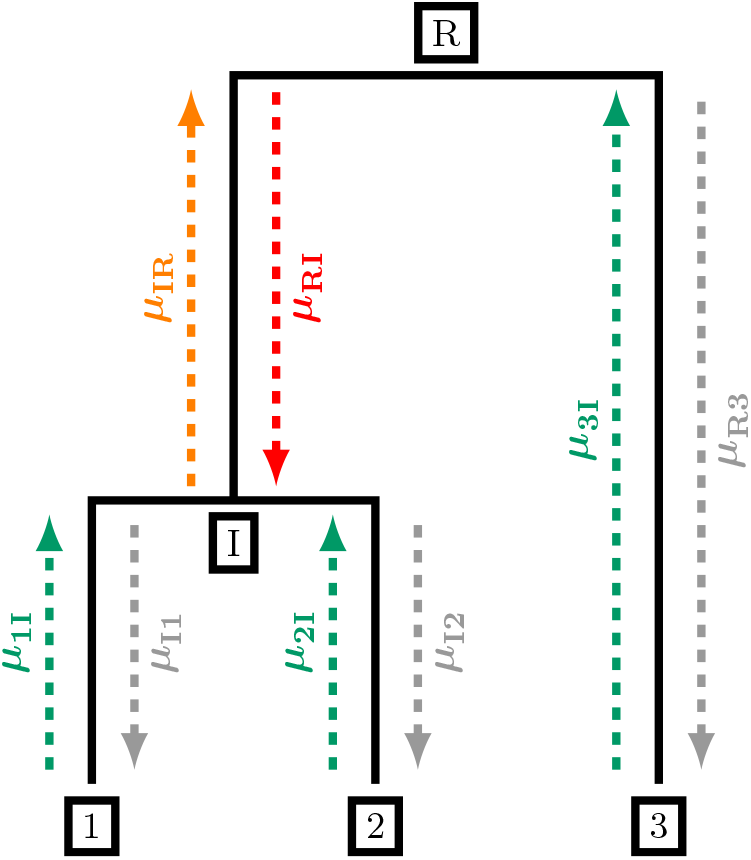
Backward filtering-forward sampling messages for a tree with 3 leaves. Messages are computed in the order green-orange-red, and messages in grey could be computed but are unnecessary to sample a new configuration of demes at nodes *I* and *R*

### Migration History Sampling

A migration history proposal is completed by sampling the migration history along each branch in the subtree conditional on the demes at the subtree leaves, root and coalescent events. Under the DTA model, the migration history along each branch is a conditionally independent realisation of a forwards-in-time Markov process given these fixed demes. We refer to these realisations of Markov processes conditioned on both their initial and final states as Markov Bridges, and there are a number of ways to sample realisations (Hobolth and Stone, 2009). We adopt the uniformization scheme in which an auxiliary Markov process is constructed with events occurring faster than in the original migration process whilst also permitting self-migrations, where a lineage migrates back into its current deme. The total number of events occurring in the auxiliary Markov process is then sampled and the deme at each jump time of the auxiliary process is sampled from a discrete-time Markov chain whilst conditioning on the deme at both ends of the branch migration history. Finally, a migration history is constructed by thinning the events and removing self-migrations in the auxiliary process.

#### Acceptance Probability

The acceptance probability of a migration history update ℋ_1_ sampled from current state ℋ_0_ is computed with the standard Metropolis–Hastings acceptance ratio

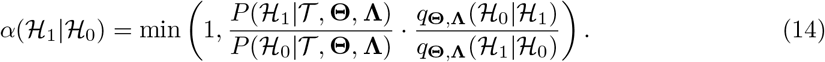

The terms in the first ratio of equation (14) are given by the structured coalescent probability density (3) evaluated for the entire structured phylogeny, noting that

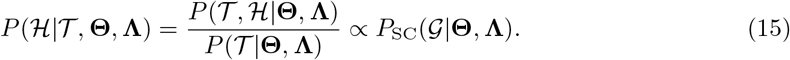

The transition kernel *q*_**Θ Λ**_ (ℋ_1_|ℋ_0_) in equation (14) can be decomposed as

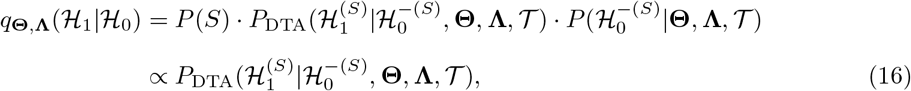

where *S* denotes the selected subtree, 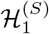 denotes the migration history of ℋ_1_ associated with *S*, and 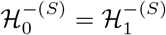 denotes the migration history of ℋ_0_ and ℋ_1_ associated with the complement of *S* respectively. This decomposition relies on the conditional independence of ℋ^(*S*)^ and ℋ^−(*S*)^ given the demes at the points reconnecting the migration history, and permits the transition kernel to evaluate the DTA probability density (4) locally on subtree *S* rather than the entire phylogeny 𝒯. Note that we evaluate the DTA probability density using forwards-in-time migration process corresponding to the current evolutionary parameters (**Θ, Λ**) using (5).

#### Implementation

The MCMC method described above is implemented into a stand-alone R package `StructCoalescent` available at https://github.com/IanPRoberts/StructCoalescent for inference using the structured coalescent on a fixed tree. Multiple utility functions, including the structured coalescent and DTA probability densities are implemented in C++ using `Rcpp` 1.0.11 (Eddelbuettel and François, 2011) and phylogenetic trees were plotted using `ape` 5.7.1 (Paradis and Schliep, 2019).

## RESULTS

### Application to a single simulated structured phylogeny

We begin by illustrating the use of our Bayesian methodology applied to a single structured phylogeny drawn from the structured coalescent probability density (3). The phylogeny was sampled using MASTER (Vaughan and Drummond, 2013), and consisted of 1000 leaves with coalescent rates sampled independently from an exponential distribution with rate 0.5 and backwards-in-time migration rates sampled independently from an exponential distribution with rate 20. Each leaf was assigned an isolation time sampled uniformly across a 50-unit time period with a sampling deme selected uniformly from the six available demes. We obtain six independent MCMC samples for this phylogeny using our default priors (9) with evolutionary parameters initialised at their respective prior means and initial migration histories sampled from the DTA probability density (4) such that each run observed a different deme at the MRCA. All runs were terminated following 48 hours of run time and completed between 2,400,000 and 2,510,000 iterations.

Our MCMC samples have mixed well over the target space with low autocorrelation in evolutionary parameter estimates and consistent regions of the target space explored in each run. Effective sample size (ESS) estimates for each evolutionary parameter exceeded 500 with mean ESS values in excess of 700, indicating relatively low autocorrelation between consecutive thinned parameter values. We also compute joint ESS (Vats et al., 2019) to assess higher-dimensional cross-correlations between groups of parameters and found estimates exceeding 1,100 for every MCMC sample. Convergence to consistent regions of the target space was assessed with the Gelman–Rubin 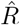 statistics (Brooks and Gelman, 1998), with every per-parameter 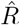 statistic below 1.002 and multivariate 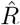 of 1.008. Whilst we are not using an explicit cutoff, such low 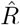 values provide no concern about a lack of convergence. Full results for ESS and 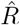 estimates of this analysis are available in Tables S1 and S2.

Convergence of evolutionary parameters alone is insufficient to conclude that MCMC samples have explored the target space effectively. We cannot directly use moment-based convergence diagnostics on samples of migration histories due to their mixed discrete and continuous components, and instead apply diagnostics to numeric summary statistics of samples of migration histories. In particular, we considered the migration frequency and total branch length observed in a deme to summarise the quantity of migration events and how they are distributed across the phylogeny. Figure 3 shows trace plots of these summary statistics for each of the six MCMC samples. Migration frequencies change frequently whilst remaining in a relatively consistent region of the posterior space in every MCMC sample (Figure 3a). We infer a median of 129 migration events (95% CI [122,139]), a little lower than the 137 migration events observed in the simulated phylogeny which we treat as a ground truth, but we maintain reasonable coverage of the ‘true’ simulation value. We see even more consistency in the stacked trace plots of branch lengths assigned to each deme (Figure 3b) with every run observing similar proportions of the migration history in each deme whilst maintaining reasonable variability at the boundaries between demes. Overall, these two sets of trace plots provide no reason to believe that we are failing to explore the space of migration histories effectively.

**Figure 3:**
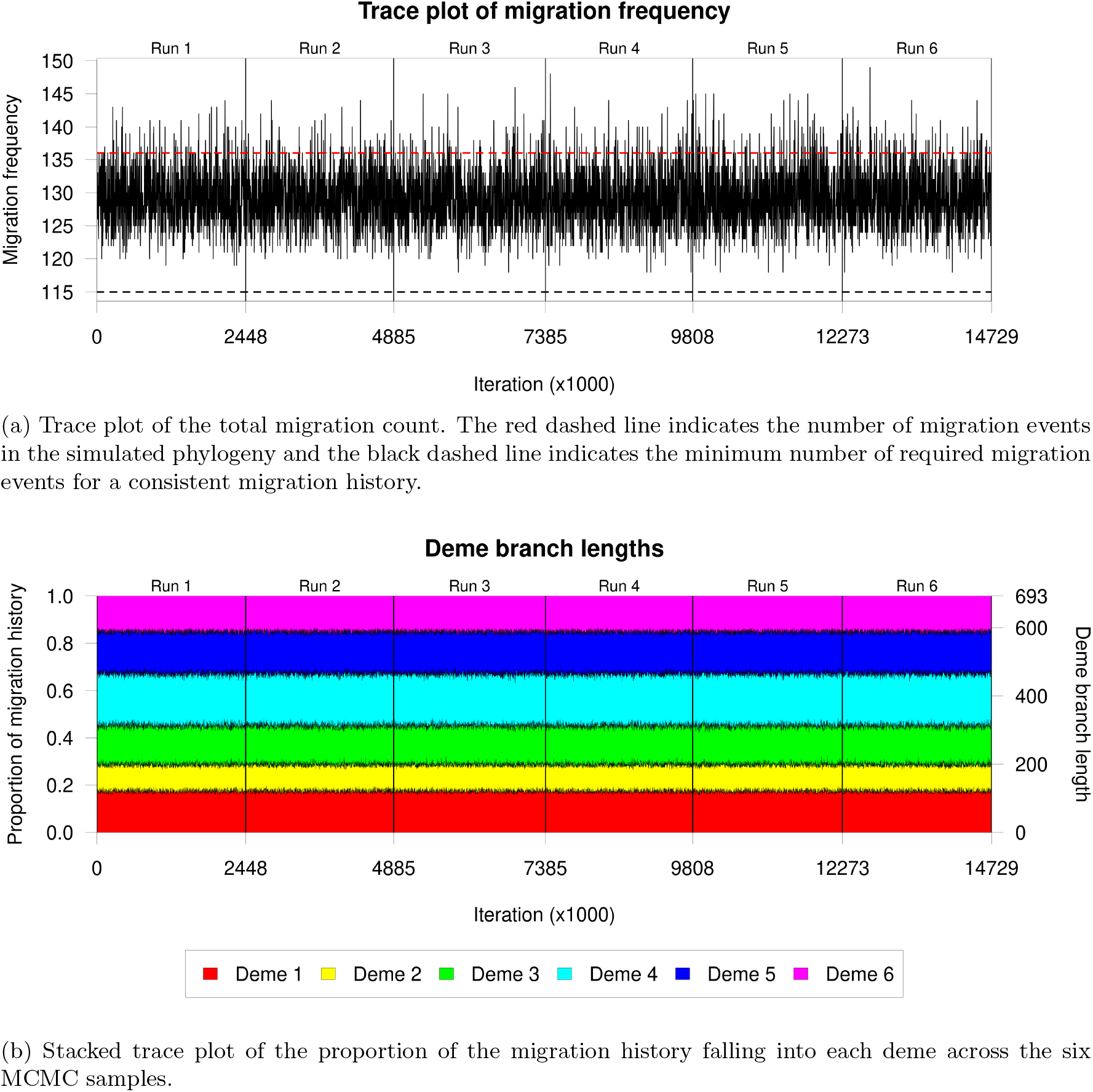

Once we are satisfied that our MCMC samples have converged over both evolutionary parameters and migration histories, we can move to assess the quality of posterior samples. We treat the evolutionary parameters used in the MASTER simulation as the ground truth and study the inferred 95% posterior credible intervals of evolutionary parameters. Simulation parameters are recovered well across the full range of values (Figure 4). Inferred coalescent rates tend to fall close to the simulation parameter in every deme with high correlation between the posterior median and simulation parameter across the full range of simulation values. We have fairly precise estimates with relatively narrow credible intervals and little systematic bias with one posterior median coalescent rate slightly underestimating the simulation parameter and one slightly overestimating the parameter. Migration rates are inferred less precisely than coalescent rates with relatively broad credible intervals and lower correlation with the posterior median, especially for simulation parameters below 0.005. This loss of precision in estimates of small migration rates has been previously noted by Müller et al. (2018) and corresponds to migration events which are rarely, if ever, observed in a sampled migration history. When no migration events are observed, the conditional posterior distributions used for Gibbs sampling are updated only by the total branch length in the source deme of the migration event. This is bounded between 0 and the total branch length of the phylogeny, although in practice we expect every deme to feature in the migration history and consequently approach this upper bound infrequently.

**Figure 4:**
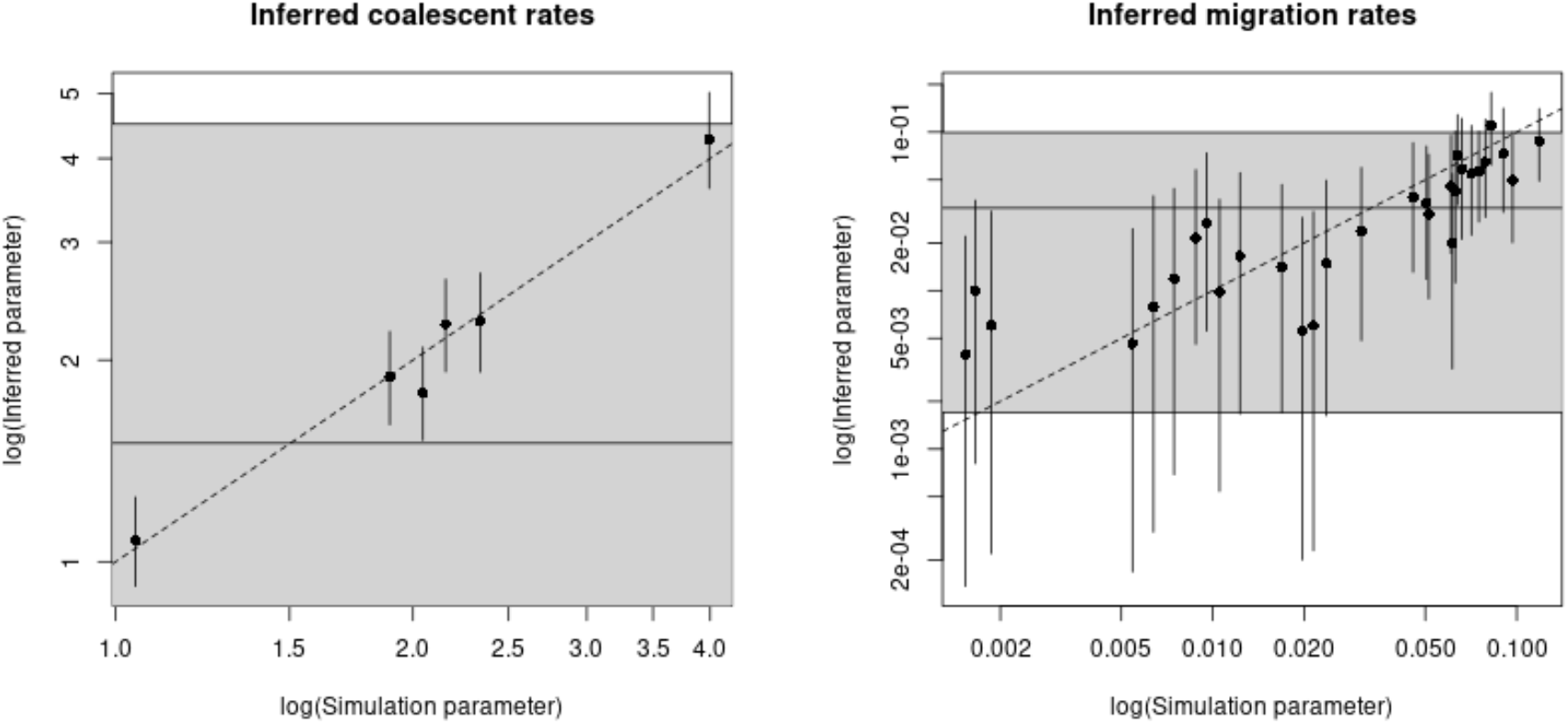
95% posterior credible intervals for coalescent rates (left) and backwards-in-time migration rates (right) aggregated over samples obtained from all six MCMC runs. The grey shaded region in each plot gives the 95% credible interval covered by the gamma-distributed priors.

We summarise MCMC samples of migration histories using a 60% majority consensus migration history (Figure 5). Each point in a majority consensus history is assigned to a deme if and only if at least a proportion *p* ∈ (0.5, 1] of sampled migration histories observe the same deme at that point. Points with insufficient consensus are assigned to a ‘null deme’ which we typically plot in black. Near the leaves, we know sampling demes with certainty and consequently there is high consensus about demes along branches close to the leaves with most of the migration history fully coloured. As we move further from the leaves, uncertainty typically increases and we find an increasing proportion of the migration history assigned to the null deme. We identify the greatest uncertainty near the MRCA with both child branches fully assigned to a null deme. The strong heterochronosity of leaf sampling and clustering of leaves within the same deme lead to fairly parsimonious migration history samples, and we identify much of the tree to be identical to the original simulation (Figure 5 inset). In the simulated migration history, we observed deme 2 in yellow at the MRCA. The pie chart at the MRCA shows that our posterior samples rarely observed deme 2 at the MRCA, with substantially lower posterior probability than any of the other five demes. We do not expect to recover every feature of the simulated migration history, and in this case it appears that the state of the simulation was relatively unlikely under our target posterior distribution.

**Figure 5:**
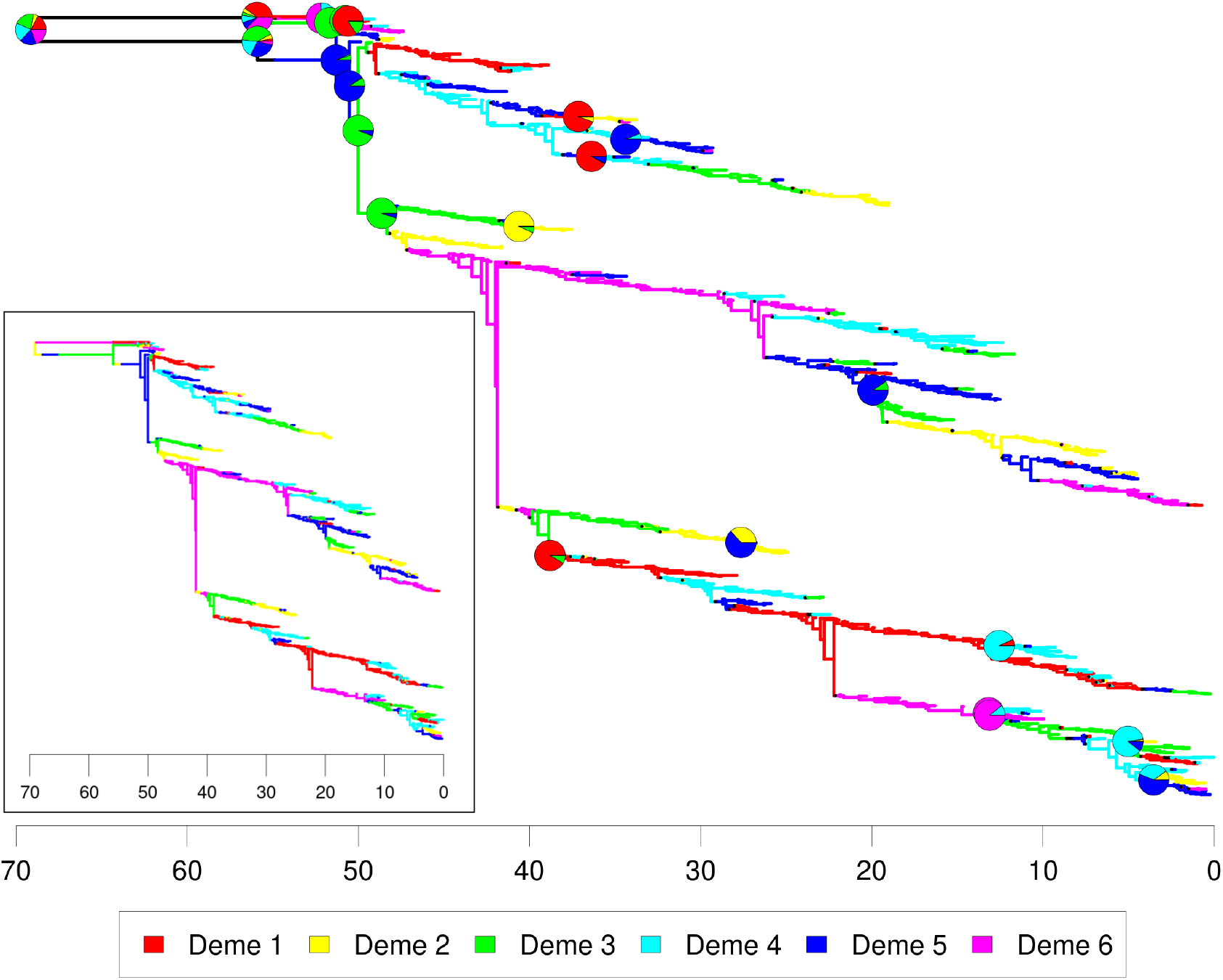
60% consensus migration history for MCMC samples on a single simulated structured phylogeny. Samples are aggregated across the six MCMC samples prior to computation and coalescent events are superimposed with pie charts denoting the posterior distribution of demes at that event. Pie charts with greater than 95% consensus are omitted, but the modal deme is indicated by the underlying migration history. Inset: Structured phylogenetic tree simulated using MASTER.

### Application to multiple simulated structured phylogenies

We completed a large-scale simulation study varying the number of demes and degree of heterochronosity. Five structured phylogenies, each with 1000 leaves were simulated using MASTER for each number of demes between 2 and 10 and four degrees of heterochronosity: lineage isolation dates sampled across a 0-unit time period (homochronous), 10-unit time period (mildly heterochronous), 30-unit time period (moderately heterochronous) or a 50-unit time period (strongly heterochronous). Coalescent rates used in simulations were sampled independently from an exponential distribution with rate 0.5 and backwards-in-time migration rates were sampled independently from an exponential distribution with rate 20. Two 48-hour MCMC samples were drawn for each phylogeny using our default prior distributions which allowed us to complete a basic convergence analysis on all 180 phylogenies.

Unlike existing methods for exact inference using the structured coalescent, we incur little penalty in per-iteration computation times with relatively stable sample sizes for all numbers of demes (Table S3). The lack of computational penalty is caused by our adaptive scheme which tends to converge to smaller subtree radii as the deme count increases. A smaller proposal radius typically reduces the number of coalescent events contained in a subtree and also reduces the branch length over which Poisson bridges must be sampled. We identify particularly slow performance for the phylogenies with 2 demes, which is also related to the adaptive scheme. Some of these phylogenies had typical acceptance probabilities in excess of our target *α*^*^ = 0.234 even when updating the entire migration history at every iteration, causing the proposal radius to increase without bound and reducing to an independence sampler. Independence sampler performance could be improved by skipping the subtree selection step, but we find these cases to be rare with more than 2 demes. A greater penalty arises from increasing the degree of heterochronosity, although this is likely caused by increasing the total size of the phylogenies with total branch lengths increasing from a mean of 176 units for homochronous phylogenies to 876 for phylogenies with strong heterochronosity.

We briefly check for signs of poor convergence in our MCMC samples using multivariate 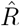 and joint ESS applied to each pair of samples (Table S3). Unfortunately, we identify multiple pairs of samples which have failed to converge to consistent distributions with multivariate 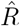 exceeding 1.2. 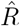 values tend to increase both as the number of demes increases, and as the degree of heterochronosity increases, with both increases indicating an increasing difficulty in sampling. Increasing the number of demes leads to a quadratic increase in the number of inferred parameters and a much more diverse space of migration histories which can make sampling challenging. Poor convergence in a single parameter is often sufficient to increase the multivariate 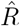 statistic to indicate poor convergence overall. The increase in 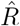 statistics associated with increasing heterochronosity may again be correlated with increases in the total branch length of the considered phylogenies. We appear to be approaching the performance limit of our Local DTA method with 48 hours of run time and phylogenies consisting of 1000 samples around 7 demes with light heterochronosity (10-units leaf sampling) although we still maintain reasonable performance with homochronous sampling up to 10 demes. Further computation time applied to these phylogenies could improve the quality of posterior samples with substantially larger sample sizes, but smaller phylogenies with fewer sampled lineages (or fewer demes) would also permit greater numbers of demes to be considered with similar run times. Considering only the 147 runs which we deem to have converged (multivariate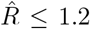), we identify very low autocorrelation in MCMC samples with almost all mean joint ESS values exceeding 1,000 and frequently exceeding 2,000 (Table S3). Joint ESS around 1,000 should be sufficient to characterise moderate quantiles of the posterior distribution although is probably insufficient for a full characterisation of events far into the tails.

We now focus on assessing the quality of evolutionary parameter samples, in particular the capacity to recover the known rates used in simulations. Figure 6 shows the inferred posterior median of each evolutionary parameter against the known simulation parameter. Both MCMC samples for each phylogeny are combined before computing posterior medians and we exclude pairs of samples with 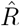 exceeding 1.2. Coalescent rates are inferred accurately across the full range of tested values, with very small deviations from the true parameter. Larger migration rates are also inferred accurately, although with less precision than coalescent rate. Small migration rates are inferred less accurately with inferred values flattening out around a posterior median of 10^−2^, again due to infrequent observation of migration events associated with the smallest migration rates.

**Figure 6:**
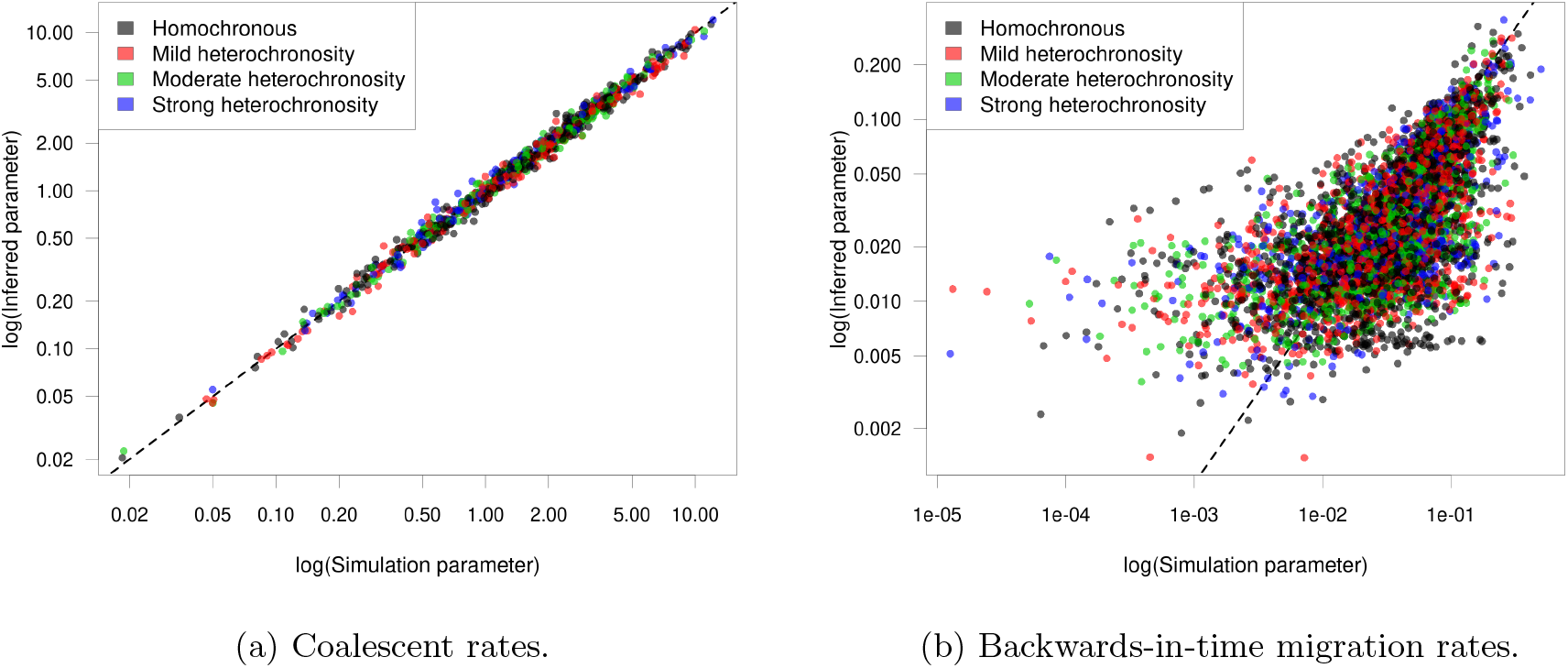
Posterior median inferred evolutionary parameter values plotted against the known simulation parameters.

We also assess the quality of inferred parameters numerically using four summary statistics:

**Coverage:** The proportion of 95% posterior credible intervals containing the simulation parameter.

**Correlation:** The correlation of the posterior median with the simulation value.

**Relative bias:** The mean deviation of the posterior samples from the simulation value normalised by the simulation value.

**Relative RMSE:** The square root of the mean squared deviation of the posterior samples from the simulation value normalised by the simulation value.

Each summary statistic is again computed by first combining pairs of MCMC samples on the same phylogeny and omitting pairs of samples with multivariate 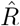 exceeding 1.2. Reported relative bias and RMSE estimates are reported only for simulation parameters greater than 5 × 10^−3^ due to the lack of statistical power in estimating small migration rates. Table 1 reports the mean of each simulation statistic taken over converged runs and separated by the degree of heterochronosity in the leaf ages and the number of demes.

**Table 1:** Coverage, correlation, relative bias and relative root mean squared error (RMSE) for inferred evolutionary parameters, separated by number of demes and degree of heterochronosity in leaf sampling. Reported values are the mean taken over pairs of MCMC samples with multivariate 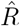 below 1.2.

|  | 2 demes | 3 demes | 4 demes | 5 demes | 6 demes | 7 demes | 8 demes | 9 demes | 10 demes |
| --- | --- | --- | --- | --- | --- | --- | --- | --- | --- |
| **Coverage** |  |  |  |  |  |  |  |  |  |
| Homochronous | 0.800 | 0.867 | 0.838 | 0.864 | 0.935 | 0.792 | 0.872 | 0.840 | 0.812 |
| Mild Heterochronosity | 0.900 | 0.956 | 0.975 | 0.910 | 0.944 | 0.918 | 0.945 | 0.922 | 0.917 |
| Moderate Heterochronosity | 0.900 | 0.926 | 0.938 | 0.912 | 0.938 | 0.939 | 0.938 | 0.914 | 0.920 |
| Strong Heterochronosity | 0.900 | 0.911 | 0.963 | 0.952 | 0.944 | 0.939 | 0.958 | 0.933 | 0.910 |
| **Posterior median correlation** |  |  |  |  |  |  |  |  |  |
| Homochronous | 0.977 | 0.999 | 0.998 | 0.997 | 0.997 | 0.996 | 0.992 | 0.993 | 0.996 |
| Mild Heterochronosity | 0.983 | 0.996 | 0.999 | 0.998 | 0.997 | 0.994 | 0.998 | 0.996 | 0.994 |
| Moderate Heterochronosity | 1.000 | 1.000 | 0.998 | 0.998 | 0.998 | 0.996 | 0.996 | 0.995 | 0.994 |
| Strong Heterochronosity | 1.000 | 0.999 | 0.998 | 0.998 | 0.998 | 0.996 | 0.996 | 0.996 | 0.992 |
| **Relative bias** |  |  |  |  |  |  |  |  |  |
| Homochronous | -0.085 | 0.581 | -0.077 | -0.048 | 0.201 | -0.099 | 0.098 | 0.057 | -0.075 |
| Mild Heterochronosity | -0.018 | 0.208 | 0.149 | 0.120 | 0.164 | 0.134 | 0.286 | 0.100 | 0.098 |
| Moderate Heterochronosity | 0.068 | 0.048 | 0.106 | 0.067 | 0.127 | 0.089 | 0.201 | 0.129 | 0.273 |
| Strong Heterochronosity | -0.004 | 0.061 | 0.183 | 0.179 | 0.116 | 0.142 | 0.180 | 0.154 | 0.141 |
| **Relative RMSE** |  |  |  |  |  |  |  |  |  |
| Homochronous | 0.499 | 1.363 | 0.812 | 0.893 | 1.107 | 0.976 | 1.159 | 1.128 | 1.040 |
| Mild Heterochronosity | 0.323 | 0.639 | 0.664 | 0.753 | 0.860 | 0.899 | 0.956 | 0.873 | 0.927 |
| Moderate Heterochronosity | 0.326 | 0.377 | 0.555 | 0.598 | 0.714 | 0.717 | 0.880 | 0.858 | 1.004 |
| Strong Heterochronosity | 0.237 | 0.423 | 0.601 | 0.634 | 0.654 | 0.755 | 0.797 | 0.839 | 0.862 |

An ideal sample from the posterior distribution should have coverage of 0.95 with the simulation parameter contained in a 95% posterior credible intervals 95% of the time. We find a mean coverage of 0.896 over the 294 accepted MCMC samples, slightly below the optimal value of 0.95 but we are still able to frequently recover the simulation parameters. The lowest coverage reliably occurs for the homochronous phylogenies (row 1) with a mean coverage of 0.847 but all other degrees of heterochronosity recover the simulation parameters more frequently. This indicates that increasing the heterochronosity within the sampled leaves provides a stronger signal to inform the inference of evolutionary parameters.

We find extremely high correlation between the posterior medians and the known simulation parameters, with all reported correlation values exceeding 0.95, indicating that estimates of the simulation parameter are close to the simulation values. Whilst this may appear inconsistent with the flattening of values observed in the plots of posterior median against simulation values (Figure 6), the observed deviations occur only for the smallest migration rates with small absolute deviations which have little effect on the correlation overall.

Due to the huge range in sizes of different evolutionary parameters, with the smallest migration rates around 10^−5^ and the largest coalescent rates greater than 10, we assess the precision of our posterior samples using relative bias and relative root mean square error (RMSE) to normalise our posterior samples. We compute the relative bias and RMSE of a sample 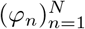 with true value *φ* as

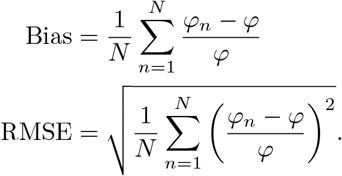

Overall, we do not find a systematic bias in our inferences with a mixture of positive and negative relative biases in our parameter estimates. Relative biases are also extremely small with few large deviations indicating accurate recovery of the simulation parameter. RMSE values are also reasonably small, indicating reasonably narrow credible intervals and fairly precise estimates of simulation parameters.

### Evolution of *Staphylococcus aureus* ST239

ST239 was the first lineage of methicillin-resistant *Staphylococcus aureus* (MRSA) to have its geographical structure investigated using whole genome sequencing (Harris et al., 2010). A phylogenetic tree from this pioneering study by Harris et al. (2010) is available on PathogenWatch (Argimón et al., 2021), consisting of 58 genomes sampled between 1982 and 2007 across 5 continents: Europe, South America, North America, Asia and Australia. This phylogeny was dated using BactDating (Didelot et al., 2018) under the additive relaxed clock model (Didelot et al., 2021). We take this dated phylogeny alongside the leaf sampling dates and locations as input and use our Local DTA methodology on this dataset to resolve the phylogeography of ST239.

In the absence of specialist knowledge about the likely migration and coalescent rates of MRSA, we again use our default priors for our analysis, in this case

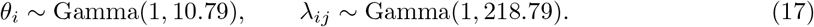

We drew five MCMC samples using these prior distributions, each terminated at 48 hours of run time, with each run completing between 3,100,000 and 3,300,000 iterations. Initial migration histories were sampled from the DTA probability density (4) such that each initial migration history sampled the MRCA in a different deme and initial coalescent and backwards-in-time migration rates were set to their respective prior means. We identify little autocorrelation within the MCMC samples with per-parameter ESS values exceeding 300 and joint ESS exceeding 1,200 in every sample. There is also no evidence of a lack of convergence with per-parameter 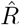 estimates below 1.005 and a multivariate 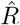 estimate of 1.023 on the basis of the five independent MCMC samples. Full results for ESS and 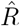 are available in figures S4, and S5 respectively.

There is no evidence to suggest that samples of migration histories have failed to converge, with reasonable exploration of migration frequencies and branch lengths in each deme (Figure S1). The range of migration counts is relatively consistent across the five MCMC samples, with some migration histories reaching maximum parsimony at a migration count of 8 and a maximum of 26 migration events observed in any run. Similarly, we find relatively consistent proportions of each deme explored whilst maintaining reasonable variability at the boundary between demes despite initialising each MCMC sample at distinct migration histories.

Our 60% majority consensus migration history (Figure 7a) is relatively parsimonious with most lineages coalescing with other lineages in the same deme before migrating away. We infer a posterior median of 13 migration events in a migration history (95% CI [9-18]), compared to a maximum parsimony migration history which requires 8 migration events. Harris et al. (2010) do not attempt to infer migration histories but note that there is strong geographical clustering between sampled MRSA lineages. Our parsimonious consensus migration history agrees well with this conclusion. We do not dine an overall consensus for the deme at the MRCA with 44% of the posterior samples originating in the Australian deme, 26% in the South American deme, 19% in the North American deme and the remaining 11% split almost evenly between Europe and Asia. The deme sampled at the MRCA may be slightly biased towards Australia due to the proximity of the only Australian sample (top-most lineage) to the MRCA. The scarcity of Australian samples provides little information about the Australian deme and we only observe the Australian lineage for a short time before the entire sample coalesces at the MRCA.

**Figure 7:**
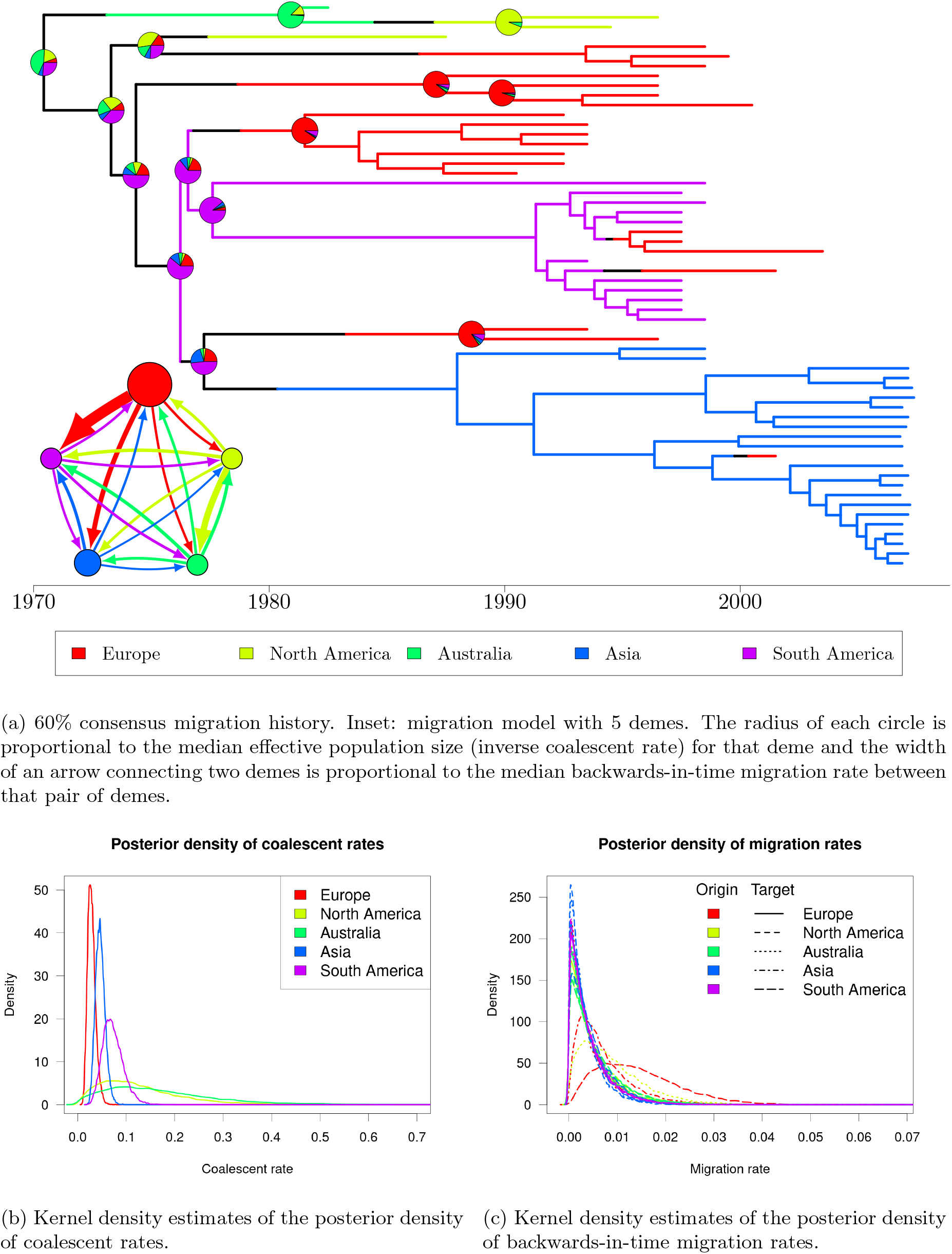
60% consensus migration history and kernel density estimates of the posterior density of evolutionary parameters for the Harris et al. (2010) MRSA analysis.

We also have access to posterior samples of coalescent and migration rates. Figures 7b and 7c shows kernel density estimates of the coalescent rates and backwards-in-time migration rates respectively. Assuming that the typical generation length of an individual remains consistent between continents and throughout time, the coalescent rate in a deme is inversely proportional to the effective population size. We observe sharp peaks in the density of coalescent rates in Europe, Asia and South America, with the respective modes indicating the smallest coalescent rates in Europe and largest in South America. From this, we can deduce that the largest MRSA population is found in Europe, with smaller population sizes in Asia and South America respectively. Despite the large population size in Europe, we have low probability of observing the European deme before 1970 in the consensus migration history, and we tend to require multiple importations of MRSA into Europe to account for its later appearance and widespread sampling. The coalescent rate estimates for the Australian and North American demes are more diffuse, with less concentrated peaks in the posterior density. This is likely due to the relative scarcity of samples from these demes - only 1 sample is available from Australia and 3 samples from North America, compared to 11 South American samples, 21 European samples and 22 Asian samples.

Almost all migration rate distributions are centred close to 0 with little distinction between different rates. The greatest inferred migration rates are the migration rates from Europe into South America and Asia, and the migration rate from North America into Australia (Figure 7a, inset), which are interpreted as infections spreading forwards-in-time from South America and Asia into Europe and from Australia into North America. The posterior distributions for the migration rate from Europe into South America is broader with a less-defined peak and consequently there is higher uncertainty in the precise value of this parameter than most of the other migration rates.

### Comparison to MultiTypeTree

MultiTypeTree (Vaughan et al., 2014) is the current state-of-the-art MCMC method for inference under the structured coalescent. It is designed to jointly infer the phylogeny, migration history and evolutionary parameters but can be adapted to inference for a fixed dated phylogeny by restricting the set of MCMC operators to the Node Retype operator to update the migration history and a Metropolis–Hastings operator to update migration rates and effective population sizes. The node retype operator is close to being a special case of our local DTA operator using a subtree consisting of the three branches connecting to a coalescent event. A coalescent event is sampled from the phylogeny at random and the migration history is updated on the two or three branches leading into or out of the coalescent event. The deme at the internal coalescent event is selected uniformly at random from the available demes and migration histories are completed using a backwards-in-time migration process. This differs from Local DTA as the node retype operator does not include information from the surrounding migration history when updating the internal deme and often has a greater probability of proposing infeasible demes which require multiple migration events to be added in order to construct a consistent migration history.

We reran our analysis of the Harris et al. (2010) MRSA phylogeny using MultiTypeTree with identical initial conditions. We used mathematically-equivalent priors to our analysis consisting of an inverse gamma prior on effective population sizes (inverse coalescent rates) and a gamma-distributed prior on backwards-in-time migration rates with the shape and rate of each prior matching our default Local DTA prior distributions. Unfortunately, we found clear evidence of a lack of convergence in every MCMC sample with divergence in both the posterior density and migration counts (Figure S2). Log-posterior densities increase rapidly from values around -300 to positive values in excess of 5000 (posterior density in excess of *e*^5000^ ≈ 10^2171^). This divergence in posterior density is likely to have been driven by the corresponding increase in migration counts, with over 1000 migration events commonly placed in a migration history.

We are unsure how or why MultiTypeTree diverged to such high migration counts - Local DTA has returned migration histories with no more than 30 migration events per migration history sampling from an identical target distribution. To eliminate the possibility of errors arising from our use of non-default prior distributions, we also reran MultiTypeTree using default lognormal(0,4) priors on both effective population sizes and migration rates. MCMC samples drawn using different prior distributions will also have different posterior distributions and are hence incomparable, but obtaining well-converged MCMC samples using lognormal priors would indicate an error in our original prior specification. We found a similar divergence to high log posterior and migration frequencies (Figure S3), although the divergence is less pronounced than using Gamma-distributed priors.

We have been unable to run MultiTypeTree stably for the Harris et al. (2010) MRSA phylogeny using two different families of prior distributions on evolutionary parameters. We are unsure what might be causing this instability but note again that MultiTypeTree is not designed with inference on a fixed phylogeny in mind, and is hence not optimised in this way.

### Determining the phylogenetic origin of Avian Influenza in Mexico

Avian influenza (AIV) is a pathogen common to a range of bird species which has been identified globally, with intercontinental spread likely to be caused by long-range migrations of wild birds (Schnebel et al., 2007). The transmission between species and across locations in Central and North America in the early 2000s was previously described based on 133 AIV genomes (Lu et al., 2014). We focus on the study of transmission between bird orders, with the 133 genomes distributed across four bird orders: Anseriformes (ANS), Charardiiformes (CHA), Galliformes (GAL) and Passeriformes (PAS), as well as transmission to domestic poultry in a more recent Mexican Outbreak (MEX). Lu et al. (2014) used a maximum likelihood method to infer a dated phylogeny relating the AIV genomes and assessed the geographic and interspecies spread using multiple sets of demes evolving as a discrete character under a DTA model. There have been at least two previous reanalyses of this dataset with De Maio et al. (2015) applying BASTA to assess interspecies spread between orders of birds, and Müller et al. (2017) applying a variety of approximations to the structured coalescent to assess the geographic spread of AIV.

We reanalyse this previously-published data from Lu et al. (2014) under the exact structured coalescent and take the dated maximum clade credibility tree inferred by Müller et al. (2017) as input. Unlike the other analyses we have carried out, we associate each AIV sample with the order of bird from which it was obtained rather than the geographic location of the sampling. Migration events are now interpreted as an interspecies transmission event rather than a geographic migration but the structured coalescent remains a reasonable model for this spread. We completed two analyses of this dataset with the first analysis consisting of five MCMC samples for which we use our default prior distributions,

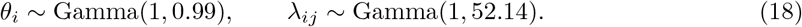

The second analysis also consisted of five MCMC samples for which we matched the previous analysis by De Maio et al. (2015) using exponentially distributed priors with rate 1. Both analyses placed almost identical priors on coalescent rates (rate 0.99 for our default priors against rate 1 for Exp(1) priors), although the migration rate priors differ substantially between the analyses. We use the same initial migration histories for both analyses with migration histories sampled from the DTA probability density (4) and initial evolutionary parameters set to the prior mean of our Local DTA priors. The initial backwards-in-time migration rate values slightly favour our first analysis using our default Gamma priors, but an Exp(1) distribution still observes high probability density at these values. Each MCMC chain was terminated at 48 hours of run time, with joint samples of evolutionary parameters saved every 100 iterations and migration histories saved every 1000 iterations.

The MCMC chains using our default priors completed a mean of 2,947,980 iterations, approximately half as many as MCMC chains using Exp(1) priors which had a mean of 5,933,560 iterations (Table 2). Despite this large discrepancy, we find that MCMC samples using Local DTA priors contain substantially lower autocorrelation with much higher ESS. Almost every MCMC sample drawn using Exp(1) priors identified at least one parameter with ESS below 100 and joint ESS values below 300. We also identified clear signs for a lack of convergence in the Exp(1) samples with a multivariate 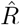 of 4.48 and 21 out of 25 evolutionary parameters with univariate 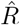 exceeding 1.1. In contrast, MCMC samples using Local DTA priors have much higher ESS values with just two parameters dropping below an ESS of 300 and every other evolutionary parameter exceeding an ESS of 500 in every run. Joint ESS was similarly much higher with every run exceeding a joint ESS of 1,000 and there were no indications of a lack of convergence with a multivariate 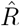 value of 1.025 and all univariate 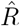 values below 1.02. Full results for ESS and 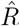 values are available in Tables S6 and S7.

**Table 2:** ESS results for evolutionary parameter estimates in the analysis of the Lu et al. (2014) AIV phylogeny using both the default prior distributions and Exp(1) prior distributions.

|  | Default priors |  |  |  |  | Exp(1) priors |  |  |  |  |
| --- | --- | --- | --- | --- | --- | --- | --- | --- | --- | --- |
|  | Run 1 | Run 2 | Run 3 | Run 4 | Run 5 | Run 1 | Run 2 | Run 3 | Run 4 | Run 5 |
| **Completed iterations** |  |  |  |  |  |  |  |  |  |  |
| $N$ | 2,931,100 | 2,965,400 | 2,925,900 | 2,965,900 | 2,951,600 | 6,870,000 | 5,909,200 | 5,408,500 | 5,308,800 | 6,171,300 |
| **Minimum ESS** |  |  |  |  |  |  |  |  |  |  |
| $\theta_{\min}$ | 928 | 1122 | 790 | 1039 | 1001 | 43 | 90 | 41 | 60 | 110 |
| $\lambda_{\min}$ | 287 | 655 | 565 | 507 | 266 | 34 | 41 | 28 | 36 | 50 |
| **Joint ESS** |  |  |  |  |  |  |  |  |  |  |
| $\Theta$ | 1095 | 1342 | 1215 | 1309 | 1183 | 98 | 258 | 108 | 170 | 108 |
| $\Lambda$ | 1208 | 1353 | 1277 | 1247 | 1232 | 130 | 179 | 201 | 164 | 137 |
| $(\Theta, \Lambda)$ | 1243 | 1446 | 1354 | 1320 | 1284 | 184 | 237 | 263 | 212 | 177 |

We also see a lack of convergence over migration histories for the Exp(1) prior MCMC samples. Trace plots of the total count of migration events in each migration history (Figure S4a) show high autocorrelation between migration histories with clear trends in the traces. We also fail to explore consistent ranges of migration counts in each run, with runs 1 and 5 typically exploring migration counts between 200 and 400, whilst runs 2, 3 and 4 typically remain between 100 and 200 migration events. There are similar issues with high autocorrelation and inconsistent regions of exploration in the stacked trace plots of branch lengths in each deme (Figure S4b). Together, these indicate that migration histories are probably mixing slowly and many more iterations would be required to achieve convergence.

Even accounting for the failure to converge, we believe that these results are sufficient evidence that Exp(1) priors are not a suitable choice for this dataset. The MCMC chains have repeatedly explored migration histories containing many more migration events than we would expect - a parsimonious reconstruction requires just 8 migration events compared with a mean of 214 sampled migration events over the five MCMC runs. We believe that this discrepancy is sufficiently large that, even if we were to run these MCMC chains for many more iterations until convergence was achieved, we would not recover biologically-reasonable results. The high number of inferred migration events is likely induced by the migration rates prior. Our default migration rates prior is also exponentially distributed with rate 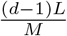, where *M* = 8 is the number of migration events required for a maximum parsimony migration history, *d* = 5 is the number of demes and *L* ≈ 104 is the total branch length in years. Reposing the Exp(1) priors in the same framework requires *M* = (*d* − 1)*L* ≈ 417, which we interpret as the prior expectation on the number of migration events under DTA. Whilst this will not be the same as the expected number of migration events under the structured coalescent, the two should be positively correlated and we can see that Exp(1) priors support much higher numbers of migration events on this phylogeny.

The large proportion of the migration history assigned to the Mexico Outbreak deme (plotted in purple) is also unlikely to be correct. Most sampled migration histories assign at least 20% of the history to this deme, with some samples extending to as much as 60% of the migration history. The Mexico H7N3 outbreak was first noted in June 2012 (Lu et al., 2014), and is unlikely to have been present and unobserved in Mexico for a long period of time prior to this date.

Returning to the results from our default Local DTA priors, we find little evidence to indicate a lack of convergence over migration histories. We infer a consistent number of migration events with most migration histories containing between 8 and 20 migration events, and have very consistent proportions of the migration history in each deme throughout all five runs (Figure S5). There is no longer a large proportion of the migration history which falls into the purple Mexico outbreak deme, with almost all migration histories containing at least 80% of the history within the red Anseriformes deme. This may be an artefact caused by biased sampling of leaves, with 93 of the 133 leaves sampled from the Anseriformes deme, but is consistent both with a parsimonious reconstruction, and the results obtained by Lu et al. (2014) using DTA.

In their analysis using Exp(1) priors and an approximation to the structured coalescent, De Maio et al. (2015) find a large amount of uncertainty about the deme at the MRCA with a reasonable probability that the MRCA may fall into any deme. De Maio et al. also use a DTA model on the same data, which recovers probability 1 of the MRCA falling into the red Anseriformes deme. We differ slightly from either of these results, with very high probability that the MRCA falls into the Anseriformes deme, but maintaining a small non-zero posterior probability of falling into any of the other four demes (Figure 8). In line with the deme branch length plots (Figure S5b), we find the majority of the migration history to be coloured in red with little variability at internal coalescent events, but observe some uncertainty in the precise position of migration events along branches and the deme at some coalescent events.

**Figure 8:**
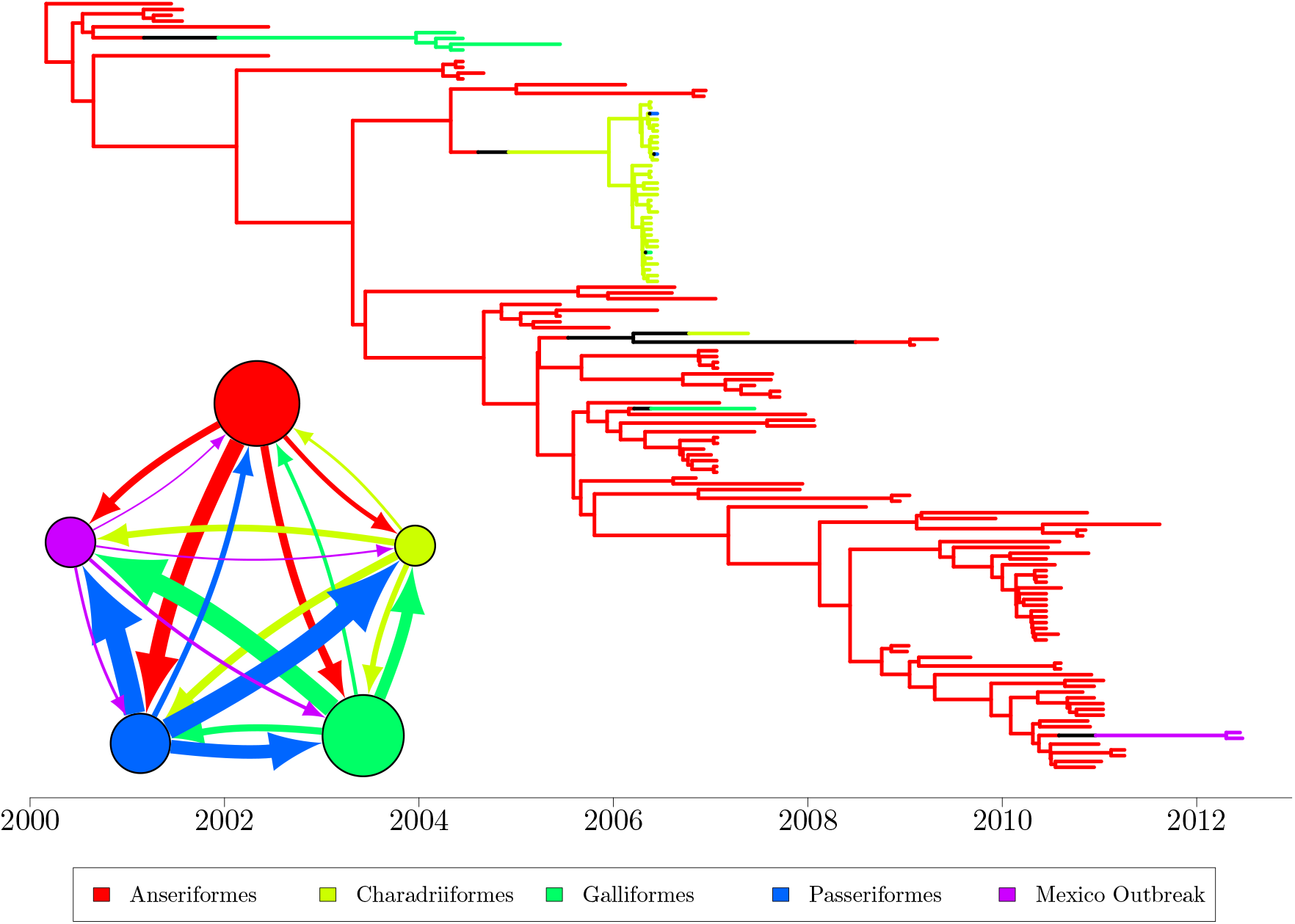
60% consensus migration history computed on all posterior migration history samples from the AIV analysis with default Local DTA priors. Inset: migration model with 5 demes. The radius of each deme circle is proportional to the median inferred effective population size (inverse coalescent rate) for that deme and the width of an arrow connecting two demes is proportional to the magnitude of the backwards-in-time migration rate between the pair of demes.

Our results with default priors bear a much greater resemblance to the results and conclusions obtained by Lu et al. (2014) and the DTA analysis by De Maio et al. (2015). We hence conclude that DTA is a reasonable approximation to the structured coalescent for this dataset when we set priors which imply relatively parsimonious reconstructions. This is likely caused by low migration rates implying that most branches contain at most one migration event irrespective of the underlying phylogeny as well as the sampling of leaves being similar to the underlying deme sizes - the greatest number of sampled lineages are drawn from the largest deme.

### Cholera Isolates from the seventh pandemic

The bacteria *Vibrio cholerae* is the causative agent of cholera, an infectious disease often thought of as only being historically important despite the fact that the seventh pandemic officially started in 1961 and is still ongoing (Baker-Austin et al., 2018). Several waves of global transmission in the seventh cholera pandemic were described on the basis of a comparison between 154 whole genomes of *V. cholerae* (Mutreja et al., 2011). A later analysis further clarified the global migration history of the seventh cholera pandemic in particular by adding many genomes from China (Didelot et al., 2015). This study was based on a total of 260 genomes sampled between 1957 and 2010 from 11 locations: Africa (AFR), Bahrain (BHR), Europe (EUR), Haiti (HTI), Indonesia (IDN), South America (SA), South Asia (SAS), Southeast Asia (SEA), China (CHN), Pakistan (PAK) and Nepal (NPL). Didelot et al. (2015) used a maximum parsimony approach (Fitch, 1971) to reconstruct a migration history, resulting in a minimum of 37 migration events, of which 18 were between an unambiguous pair of demes. The greatest number of migrations were found to have occurred from South Asia into China, and vice versa.

We reanalysed this previously-published data from Didelot et al. (2015) using the structured coalescent, adding explicit sampling of migration histories to the original analysis. We take as input the same dated phylogeny (Figure S6) and draw eleven MCMC samples with initial migration histories sampled from the DTA probability density such that each run observes a unique deme at the MRCA. Our preliminary analysis using our default Gamma-distributed priors failed to fully converge with some sharp changes in migration count throughout the MCMC samples (Figure S7. The migration histories also frequently featured very high prevalence of the Haiti deme with many migration histories observing at least 10% of the history occurring in Haiti (Figure S8). This requires many migration events into and out of Haiti as well as contradicting the scientific consensus that seventh pandemic Cholera entered Haiti in early 2010 following at least a century since the previous Cholera case (Chin et al., 2011; Hendriksen et al., 2011). In light of these issues, we repeated our analysis using a more conservative prior on migration rates to reduce the prior expected number of migration events. We found that this resulted in more parsimonious migration histories and we were able to identify some known events in the spread of seventh pandemic Cholera.

The rate of our default prior on migration rates (9) is specified using the number of demes *d*, total branch length of the phylogeny ℒ and a prior belief about the number of migration events *M*. By default, we set *M* to be the number of required migration events in a maximum parsimony migration history, in this case *M* = 37 migration events. We approximately halved *M* to a value of *M* = 20, resulting in reduced prior distributions

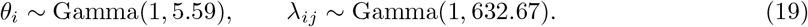

Each of the eleven MCMC runs completed around 2,500,000 iterations and obtain reasonably high quality samples. Every evolutionary parameter sample contained reasonably low autocorrelation with ESS values of at least 250 and mean per-parameter ESS of 1,921. Joint ESS values were similarly high with a minimum joint ESS of 2,901. All samples also converged well to a consistent distribution with 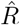 statistics of at most 1.03. Full results for ESS and 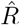 are available in Tables S9 and S10.

We also find no evidence to indicate poor convergence over migration histories. Sampled migration histories tend to observe between 45 and 50 migration events with migration histories occasionally dropping to 37 migration events, matching with the number of migration histories required for a maximum parsimony migration history (Figure S9a). We also have extremely consistent proportions of the migration history assigned to each deme, and notably observe very small proportions of the Haitian deme unlike our preliminary analysis (Figure S9b).

Figure 9 shows a 60% consensus migration history with the median inferred effective population sizes and backwards-in-time migration rates shown inset. We find very high probability of the MRCA arising in Asia with 92% of the sampled migration histories identifying the MRCA in Indonesia, 4.8% in South East Asia, 1.8% in China and the remaining 1.4% split between the remaining demes. There remains some uncertainty about the precise origin within Asia, but the inference is likely to depend on the sampling scheme for lineages in the phylogeny. Most of the lineages which are isolated close to the MRCA are obtained either from China or Indonesia, and the resulting migration history would require a strong migration process to migrate away from either deme in such a short period of time.

**Figure 9:**
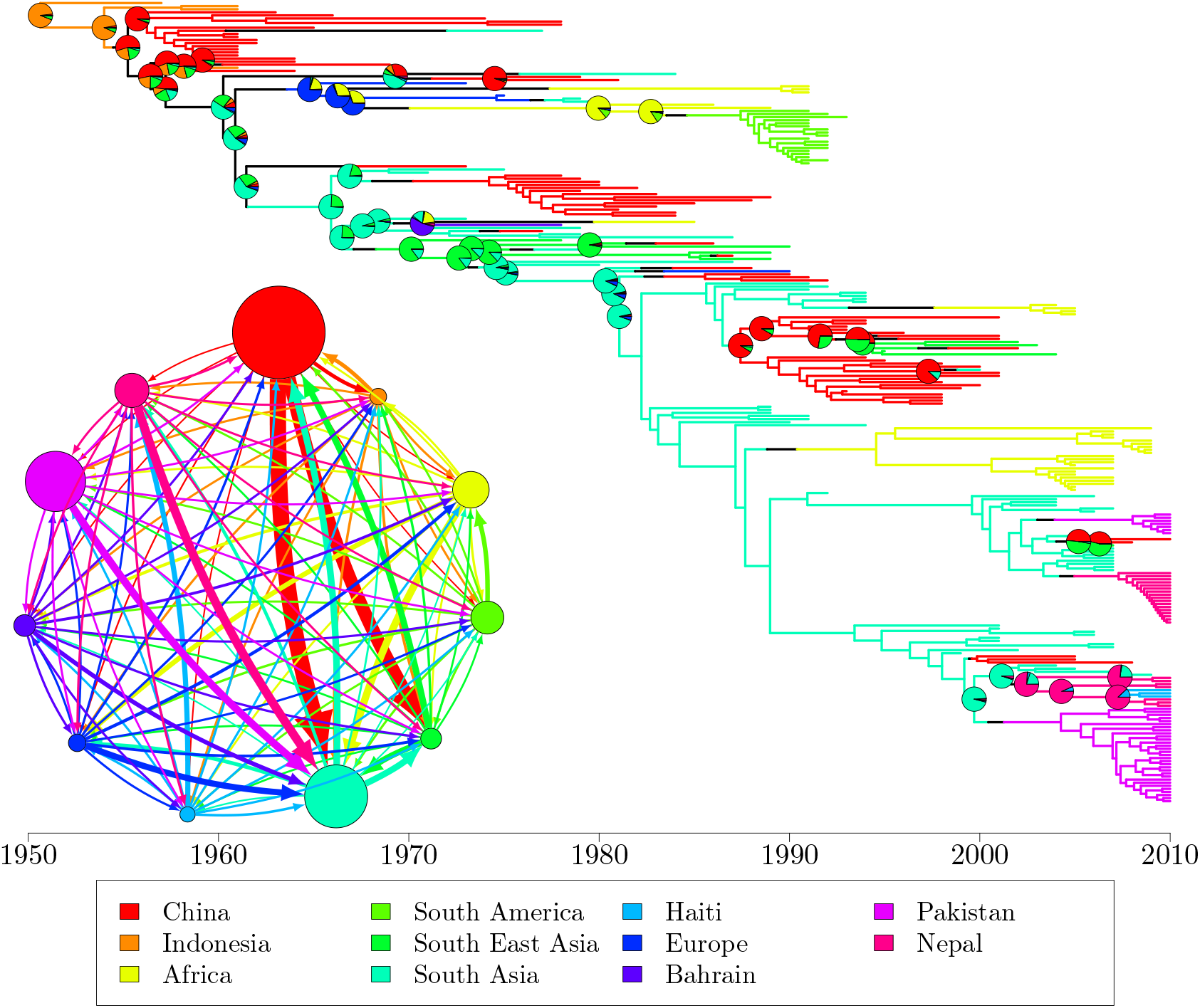
60% consensus migration history based on all migration histories sampled across the eleven MCMC samples in our Cholera analysis. Inset: migration model with 11 demes. The radius of each deme circle is proportional to the median inferred effective population size for that deme and the width of an arrow connecting two demes is proportional to the magnitude of the backwards-in-time migration rate between the pair of demes.

We now focus on two well-documented transmission events which are recovered by our analysis, beginning with the introduction of seventh pandemic Cholera into Haiti. Cholera is widely believed to have been reintroduced into Haiti following an earthquake in early 2010 by Nepalese aid workers following at least a century since the previous Haitian cholera outbreak (Chin et al., 2011; Hendriksen et al., 2011). Figure S10a shows the migration history focused around the three Haitian Cholera isolates. We infer the transmission event into Haiti to occur between 2007 and 2008, between 2 and 3 years earlier than would be expected. We believe that the early appearance of this event is likely caused by an explosive population growth of Cholera within Haiti (forwards-in-time) following introduction. There was no endemic population of seventh pandemic Cholera strains within Haiti prior to transmission from Nepal and we would consequently expect rapid growth after introduction. This violates the assumption of fixed deme sizes (equivalently fixed coalescent rates) under the structured coalescent and makes precise inference in this region of the phylogeny unreliable.

The other transmission event we mention is the introduction of seventh pandemic Cholera into South America. Cholera was identified in South America in January 1991 (Centers for Disease Control and Prevention, 1991; Guthmann, 1995), although the origin of the transmission was initially unclear. It was originally suggested that Cholera was reintroduced into South America from Asia by international trade ships, but more recent work has found genetic similarities between the African and South American strains which are not present in Asian samples (Lam et al., 2010). We agree with the more recent work and infer a transmission of Cholera from Africa into South America between 1983 and 1986. Much like our inference of transmission into Haiti, we infer the transmission event between 5 and 8 years earlier than the true transmission event although this is likely caused by a similar rapid growth immediately following introduction.

We identify China as the largest population of seventh pandemic Cholera with other large populations in South Asia and Pakistan (Figure 9, inset). We also find the greatest backwards-in-time migration rates to be those from China into South Asia and South East Asia, with most of the largest backwards-in-time migration rates targeting South East Asia. We have less capacity to distinguish between the smallest inferred migration rates and instead consider the number of migration events backwards-in-time (Figure S11). The number of migration events between a pair of deme is usually positively correlated with the migration rate with higher migration rates usually inducing larger quantities of migration events. If many samples are drawn from the same deme, we may still observe many migration events even with low migration rates, especially if the samples are separated into multiple sampling groups. In line with Didelot et al. (2015)’s findings, we identify the greatest number of migration events between China, South Asia and Southeast Asia with relatively few migration events between other pairs of demes.

## DISCUSSION

We have introduced a modular Bayesian inference framework for inference using the structured coalescent and contribute a Bayesian approach ‘Local DTA’ for inference with the structured coalescent conditional on a dated phylogenetic tree. Our inferential framework requires a dated phylogenetic tree to be first estimated from sequence data, for which there are many previously-published methods. There are no existing methods explicitly designed for the second step moving from an unstructured dated phylogeny to full structured phylogenies, although two existing MCMC schemes (Ewing et al., 2004; Vaughan et al., 2014) may be adapted for this inference by restricting the set of MCMC operators.

We have applied Local DTA to a range of simulated and previously-published empirical datasets. In simulations, we show clear evidence that Local DTA converges well for phylogenetic trees containing 1000 leaves and up to 10 demes with reasonable computation times up to 48 hours. We recover known simulation parameters well with highly accurate and precise estimates of known parameters for all but the smallest migration rates, which have been previously-noted to be poorly informed in phylogeographic analyses (Müller et al., 2017). Applied to previously-published MRSA, AIV and Cholera datasets, we were able to replicate previously-described conclusions about the ancestral structure of sampled lineages. In particular, we inferred two well-known transmission events in the seventh Cholera pandemic with high probability, although our inferred transmission times were somewhat earlier than the true events. Bayesian inference using the structured coalescent is highly prior dependent, typically with high correlation between the number of inferred migration events and the expectation of migration rate priors. The AIV analysis illustrates this dependence with a comparison between our default prior distributions and a previously-described analysis using less conservative exponential priors (De Maio et al., 2015). The less conservative estimate showed clear signs of a lack of convergence as well as supporting many more migration events than would be expected.

