## Supplementary Material for "Bayesian Inference of Pathogen Phylogeography using the Structured Coalescent Model"

| | Run 1 | Run 2 | Run 3 | Run 4 | Run 5 | Run 6 | **Min** | **Max** | **Mean** |
|---|---|---|---|---|---|---|---|---|---|
| **Coalescent rates** | | | | | | | | | |
| $\theta_1$ | 1110 | 979 | 871 | 896 | 934 | 992 | **871** | **1110** | **964** |
| $\theta_2$ | 1284 | 1119 | 1401 | 956 | 929 | 1306 | **929** | **1401** | **1166** |
| $\theta_3$ | 781 | 775 | 1047 | 974 | 1004 | 823 | **775** | **1047** | **901** |
| $\theta_4$ | 1156 | 1154 | 987 | 970 | 909 | 1036 | **909** | **1156** | **1035** |
| $\theta_5$ | 927 | 1270 | 755 | 778 | 962 | 1036 | **755** | **1270** | **955** |
| $\theta_6$ | 858 | 1025 | 991 | 1445 | 1185 | 1399 | **858** | **1445** | **1150** |
| **Migration rates** | | | | | | | | | |
| $\lambda_{2,1}$ | 953 | 1024 | 1257 | 824 | 997 | 935 | **824** | **1257** | **998** |
| $\lambda_{3,1}$ | 1004 | 841 | 927 | 907 | 1293 | 757 | **757** | **1293** | **955** |
| $\lambda_{4,1}$ | 1153 | 849 | 759 | 795 | 1077 | 832 | **759** | **1153** | **911** |
| $\lambda_{5,1}$ | 1234 | 1015 | 726 | 849 | 949 | 871 | **726** | **1234** | **941** |
| $\lambda_{6,1}$ | 852 | 1003 | 830 | 964 | 1043 | 729 | **729** | **1043** | **904** |
| $\lambda_{1,2}$ | 856 | 1084 | 967 | 810 | 963 | 1269 | **810** | **1269** | **992** |
| $\lambda_{3,2}$ | 794 | 1158 | 704 | 728 | 1127 | 1033 | **704** | **1158** | **924** |
| $\lambda_{4,2}$ | 1360 | 1170 | 865 | 1334 | 1184 | 1079 | **865** | **1360** | **1165** |
| $\lambda_{5,2}$ | 918 | 686 | 832 | 879 | 1153 | 829 | **686** | **1153** | **883** |
| $\lambda_{6,2}$ | 1123 | 897 | 837 | 755 | 1051 | 1217 | **755** | **1217** | **980** |
| $\lambda_{1,3}$ | 807 | 971 | 835 | 1008 | 942 | 775 | **775** | **1008** | **890** |
| $\lambda_{2,3}$ | 937 | 899 | 1013 | 1074 | 923 | 939 | **899** | **1074** | **964** |
| $\lambda_{4,3}$ | 822 | 740 | 1024 | 850 | 1144 | 1172 | **740** | **1172** | **959** |
| $\lambda_{5,3}$ | 957 | 789 | 783 | 1570 | 960 | 1365 | **783** | **1570** | **1071** |
| $\lambda_{6,3}$ | 873 | 808 | 836 | 692 | 1254 | 824 | **692** | **1254** | **881** |
| $\lambda_{1,4}$ | 549 | 763 | 844 | 773 | 746 | 659 | **549** | **844** | **722** |
| $\lambda_{2,4}$ | 1067 | 773 | 870 | 893 | 1048 | 885 | **773** | **1067** | **923** |
| $\lambda_{3,4}$ | 843 | 785 | 968 | 781 | 761 | 1015 | **761** | **1015** | **859** |
| $\lambda_{5,4}$ | 745 | 841 | 915 | 687 | 998 | 851 | **687** | **998** | **840** |
| $\lambda_{6,4}$ | 704 | 741 | 736 | 1002 | 885 | 763 | **704** | **1002** | **805** |
| $\lambda_{1,5}$ | 832 | 1143 | 905 | 828 | 1148 | 808 | **808** | **1148** | **944** |
| $\lambda_{2,5}$ | 850 | 795 | 721 | 790 | 999 | 826 | **721** | **999** | **830** |
| $\lambda_{3,5}$ | 742 | 690 | 783 | 1159 | 1049 | 1084 | **690** | **1159** | **918** |
| $\lambda_{4,5}$ | 853 | 842 | 1057 | 760 | 820 | 871 | **760** | **1057** | **867** |
| $\lambda_{6,5}$ | 865 | 918 | 858 | 807 | 1208 | 811 | **807** | **1208** | **911** |
| $\lambda_{1,6}$ | 737 | 682 | 816 | 923 | 932 | 820 | **682** | **932** | **818** |
| $\lambda_{2,6}$ | 1045 | 1030 | 899 | 817 | 1039 | 1074 | **817** | **1074** | **984** |
| $\lambda_{3,6}$ | 1113 | 965 | 1077 | 971 | 1098 | 1115 | **965** | **1115** | **1056** |
| $\lambda_{4,6}$ | 985 | 922 | 822 | 934 | 999 | 1058 | **822** | **1058** | **953** |
| $\lambda_{5,6}$ | 914 | 779 | 941 | 783 | 1003 | 842 | **779** | **1003** | **877** |

Table S1: Effective sample sizes of evolutionary parameters for simulation study on a single structured phylogeny, computed in R using `mcmcse` (Flegal et al., 2021).

| | $\theta_x$ | $\lambda_{x,1}$ | $\lambda_{x,2}$ | $\lambda_{x,3}$ | $\lambda_{x,4}$ | $\lambda_{x,5}$ | $\lambda_{x,6}$ |
|---|---|---|---|---|---|---|---|
| $x = 1$ | 1.00076 | — | 1.00020 | 1.00067 | 1.00076 | 1.00114 | 1.00064 |
| $x = 2$ | 1.00071 | 1.00030 | — | 1.00094 | 1.00038 | 1.00091 | 1.00080 |
| $x = 3$ | **1.00096** | 1.00074 | 1.00334 | — | 1.00032 | 1.00011 | 1.00091 |
| $x = 4$ | 1.00050 | 1.00128 | 1.00036 | 1.00095 | — | 1.00099 | 1.00119 |
| $x = 5$ | 1.00068 | **1.00191** | 1.00040 | 1.00086 | 1.00089 | — | 1.00120 |
| $x = 6$ | 1.00062 | 1.00101 | 1.00111 | 1.00144 | 1.00197 | 1.00039 | — |

Table S2: Gelman–Rubin $\hat{R}$ values for the evolutionary parameters, computed using the `coda` R package (Plummer et al., 2006), with the greatest $\hat{R}$ value highlighed in **bold**. The second column gives the $\hat{R}$ value for the coalescent rate in each deme whilst the remaining columns give the $\hat{R}$ values for the backwards-in-time migration rates. The row of the table gives the source deme of each migration rate, and the column gives the target deme.

| | 2 demes | 3 demes | 4 demes | 5 demes | 6 demes | 7 demes | 8 demes | 9 demes | 10 demes |
|---|---|---|---|---|---|---|---|---|---|
| **Completed iterations** ($\times 1000$) | | | | | | | | | |
| Homochronous | 923 | 1,869 | 3,386 | 4,082 | 4,750 | 4,809 | 4,580 | 3,938 | 5,207 |
| Mild Heterochronosity | 925 | 2,537 | 3,015 | 2,789 | 4,742 | 2,872 | 3,611 | 4,403 | 4,930 |
| Moderate Heterochronosity | 1,517 | 3,190 | 3,689 | 2,841 | 3,337 | 2,849 | 3,405 | 3,001 | 3,327 |
| Strong Heterochronosity | 1,846 | 3,007 | 2,889 | 3,135 | 3,500 | 3,678 | 3,589 | 3,788 | 3,867 |
| **Multivariate** $\hat{R} \leq 1.2$ | | | | | | | | | |
| Homochronous | 5 | 5 | 5 | 5 | 3 | 5 | 5 | 4 | 5 |
| Mild Heterochronosity | 5 | 5 | 5 | 4 | 5 | 2 | 2 | 3 | 3 |
| Moderate Heterochronosity | 5 | 3 | 5 | 5 | 4 | 3 | 5 | 1 | 1 |
| Strong Heterochronosity | 5 | 5 | 5 | 5 | 5 | 4 | 3 | 5 | 2 |
| **Joint ESS** | | | | | | | | | |
| Homochronous | 819 | 1,564 | 2,403 | 2,980 | 2,392 | 3,145 | 2,811 | 3,263 | 3,122 |
| Mild Heterochronosity | 791 | 2,020 | 2,323 | 2,187 | 2,437 | 2,616 | 2,103 | 2,061 | 2,165 |
| Moderate Heterochronosity | 1,257 | 1,786 | 2,092 | 1,813 | 2,100 | 2,211 | 1,736 | 1,436 | 1,677 |
| Strong Heterochronosity | 1,621 | 2,033 | 1,975 | 1,989 | 1,683 | 1,611 | 1,723 | 1,551 | 1,341 |

Table S3: Summary of convergence diagnostics for inferred evolutionary parameters, separated by number of demes and degree of heterochronosity in leaf sampling. Reported numbers of iterations and joint ESS values are means taken over all MCMC samples of that type.

| | Run 1 | Run 2 | Run 3 | Run 4 | Run 5 | Run 6 | Run 7 | Run 8 | Run 9 | Run 10 | Run 11 | Run 12 |
|---|---|---|---|---|---|---|---|---|---|---|---|---|
| **Coalescent rates** | | | | | | | | | | | | |
| $\theta_{\mathrm{EUR}}$ | 924.50 | 1236.94 | 1195.81 | 1032.50 | **866.00** | 956.01 | 922.59 | 1335.68 | 1109.84 | 1172.58 | 1012.66 | 1103.95 |
| $\theta_{\mathrm{NA}}$ | 1119.75 | 1148.91 | 1434.47 | 932.17 | 1128.25 | **903.11** | 931.58 | 1529.84 | 1253.96 | 1501.70 | 1254.16 | 1103.64 |
| $\theta_{\mathrm{AUS}}$ | 1056.95 | 917.31 | 968.27 | 815.52 | 1391.40 | 1093.73 | 1047.46 | **798.97** | 1062.77 | 1224.33 | 1541.31 | 1198.59 |
| $\theta_{\mathrm{AS}}$ | 1435.85 | **1117.87** | 1559.84 | 1288.67 | 1277.33 | 1263.61 | 1400.34 | 1287.97 | 1689.01 | 1429.37 | 1525.23 | 1185.07 |
| $\theta_{\mathrm{SA}}$ | 1050.64 | 897.98 | 974.62 | 1128.48 | 1000.09 | 979.07 | 944.11 | 1163.00 | **826.60** | 1010.93 | 1100.99 | 1034.01 |
| **Migration rates** | | | | | | | | | | | | |
| $\lambda_{\mathrm{NA,EUR}}$ | 1047.23 | 937.85 | 946.02 | 1033.56 | **932.81** | 1103.67 | 1258.02 | 1112.22 | 1282.76 | 1262.66 | 1029.33 | 1126.62 |
| $\lambda_{\mathrm{AUS,EUR}}$ | 1695.76 | 1195.74 | 1173.14 | 1231.09 | 1066.31 | 1062.40 | 1555.46 | 1952.03 | 1580.97 | 1214.71 | **1026.67** | 1423.33 |
| $\lambda_{\mathrm{AS,EUR}}$ | 987.94 | 963.85 | 908.26 | 1163.59 | 1075.44 | **894.86** | 1296.94 | 1234.26 | 1249.47 | 902.20 | 1137.79 | 1134.13 |
| $\lambda_{\mathrm{SA,EUR}}$ | 1159.55 | 808.07 | 1033.64 | 857.67 | 883.37 | 873.25 | 941.90 | **708.89** | 1073.21 | 1055.83 | 1023.25 | 897.18 |
| $\lambda_{\mathrm{EUR,NA}}$ | 807.27 | 778.51 | 1079.50 | 919.19 | 618.13 | **610.17** | 819.34 | 823.20 | 1196.97 | 710.62 | 940.65 | 796.41 |
| $\lambda_{\mathrm{AUS,NA}}$ | 1248.54 | 1284.42 | 1251.94 | 1037.22 | 1018.41 | 1197.83 | 903.47 | **710.22** | 1291.43 | 1114.64 | 947.01 | 728.39 |
| $\lambda_{\mathrm{AS,NA}}$ | 1323.98 | 890.10 | 1047.13 | 1049.42 | **875.12** | 1077.93 | 926.02 | 1401.17 | 1320.30 | 1111.81 | 1225.77 | 1472.49 |
| $\lambda_{\mathrm{SA,NA}}$ | 1046.14 | 1111.14 | 1210.28 | 931.84 | 1402.89 | 1235.25 | **768.47** | 1073.22 | 930.91 | 1132.29 | 909.55 | 847.63 |
| $\lambda_{\mathrm{EUR,AUS}}$ | 378.18 | 747.75 | 473.86 | 536.90 | **314.85** | 536.74 | 784.07 | 574.83 | 365.99 | 368.33 | 430.63 | 511.26 |
| $\lambda_{\mathrm{NA,AUS}}$ | 1016.55 | 1172.97 | 1204.82 | 1193.43 | 1243.00 | **805.79** | 1234.99 | 1048.86 | 1055.11 | 1251.52 | 833.46 | 1415.35 |
| $\lambda_{\mathrm{AS,AUS}}$ | 492.45 | **466.41** | 1707.95 | 1163.83 | 854.62 | 1011.09 | 1042.03 | 1482.01 | 894.61 | 1028.70 | 1116.78 | 1527.67 |
| $\lambda_{\mathrm{SA,AUS}}$ | 1070.80 | 1178.21 | 1033.25 | 853.98 | 1083.96 | 1221.74 | 1082.33 | 891.26 | **668.35** | 1285.66 | 1065.27 | 1206.00 |
| $\lambda_{\mathrm{EUR,AS}}$ | 713.63 | 543.13 | 630.52 | 655.21 | 510.78 | 622.36 | **504.51** | 588.68 | 694.94 | 670.33 | 748.08 | 593.74 |
| $\lambda_{\mathrm{NA,AS}}$ | 1259.62 | **879.92** | 1324.68 | 1619.02 | 1027.18 | 1232.26 | 1152.38 | 1302.36 | 1397.13 | 1408.76 | 1293.60 | 979.71 |
| $\lambda_{\mathrm{AUS,AS}}$ | 1598.62 | 1278.63 | 1396.77 | 1067.92 | 1352.89 | **802.16** | 1138.57 | 999.72 | 1293.07 | 1372.60 | 1236.75 | 1251.17 |
| $\lambda_{\mathrm{SA,AS}}$ | 1085.89 | 833.83 | 977.34 | 963.61 | 760.64 | **677.21** | 901.40 | 841.37 | 985.64 | 1018.01 | 1044.32 | 778.65 |
| $\lambda_{\mathrm{EUR,SA}}$ | 631.94 | 569.84 | 623.95 | 564.81 | 477.66 | 442.44 | 675.13 | 638.14 | 589.67 | **437.14** | 442.23 | 601.04 |
| $\lambda_{\mathrm{NA,SA}}$ | 976.04 | 1156.54 | 1032.97 | 920.27 | 1023.72 | 1304.43 | 1136.74 | 1113.00 | 884.55 | 1171.80 | **668.57** | 982.67 |
| $\lambda_{\mathrm{AUS,SA}}$ | 1215.03 | 1036.15 | 1065.80 | 1203.26 | 1002.00 | 1318.34 | **920.75** | 1209.34 | 1299.39 | 1180.04 | 1036.35 | 1058.19 |
| $\lambda_{\mathrm{AS,SA}}$ | 898.71 | **728.15** | 1071.80 | 757.15 | 961.99 | 1207.40 | 1094.43 | 1047.46 | 1168.38 | 1085.08 | 745.50 | 1031.66 |

Table S4: Effective sample size estimates for each evolutionary parameter in the MRSA analysis.

| | $\lambda_{\mathrm{EUR},x}$ | $\lambda_{\mathrm{NA},x}$ | $\lambda_{\mathrm{AUS},x}$ | $\lambda_{\mathrm{AS},x}$ | $\lambda_{\mathrm{SA},x}$ | $\theta_x$ |
|---|---|---|---|---|---|---|
| $x$=EUR | — | 1.0016 (1.0028) | 1.0034 (1.0051) | 1.0006 (1.0010) | 1.0012 (1.0022) | 1.0008 (1.0015) |
| $x$=NA | 1.0005 (1.0008) | — | 1.0008 (1.0015) | 1.0008 (1.0014) | 1.0006 (1.0010) | 1.0004 (1.0006) |
| $x$=AUS | 1.0009 (1.0015) | 1.0005 (1.0008) | — | 1.0007 (1.0012) | 1.0007 (1.0012) | 1.0006 (1.0010) |
| $x$=AS | 1.0005 (1.0009) | 1.0008 (1.0014) | 1.0010 (1.0015) | — | 1.0005 (1.0009) | 1.0005 (1.0010) |
| $x$=SA | 1.0009 (1.0015) | 1.0007 (1.0012) | 1.0008 (1.0012) | 1.0008 (1.0012) | — | 1.0008 (1.0016) |

Table S5: Gelman-Rubin $\hat{R}$ estimates for each evolutionary parameter in the MRSA analysis. Quantities in brackets give the upper 95% confidence interval for the $\hat{R}$ value.

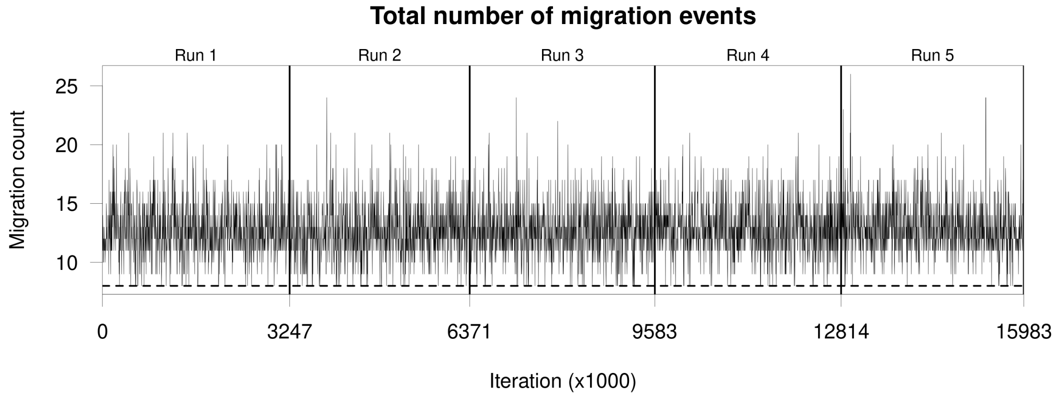


(a) Trace plot of total migration counts. The dashed line at a migration count of 8 indicates the minimum required number of migration events for a maximum parsimony migration history.

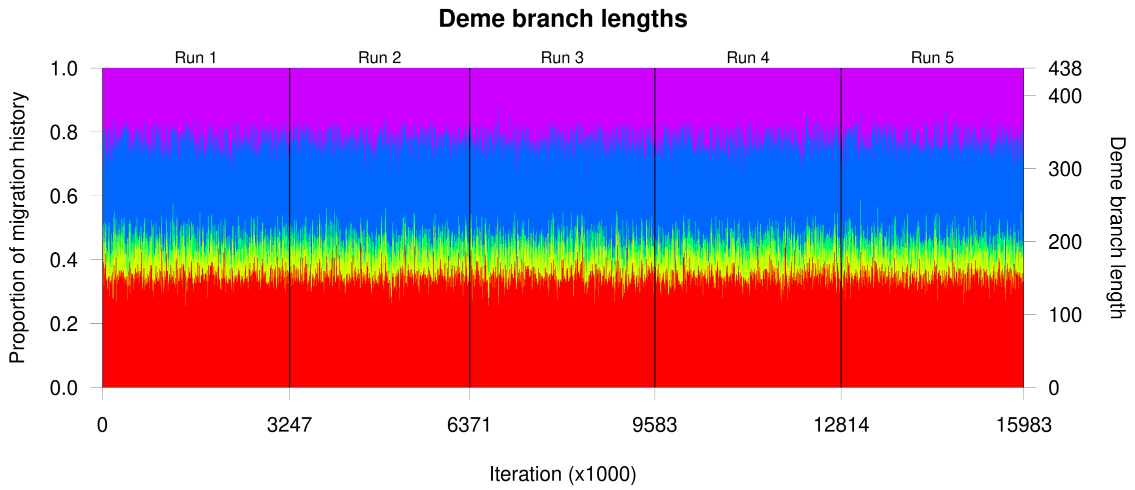


(b) Stacked trace plot of total deme branch lengths

Figure S1: Summary statistics for mixing over migration histories for the twelve runs in the MRSA analysis.

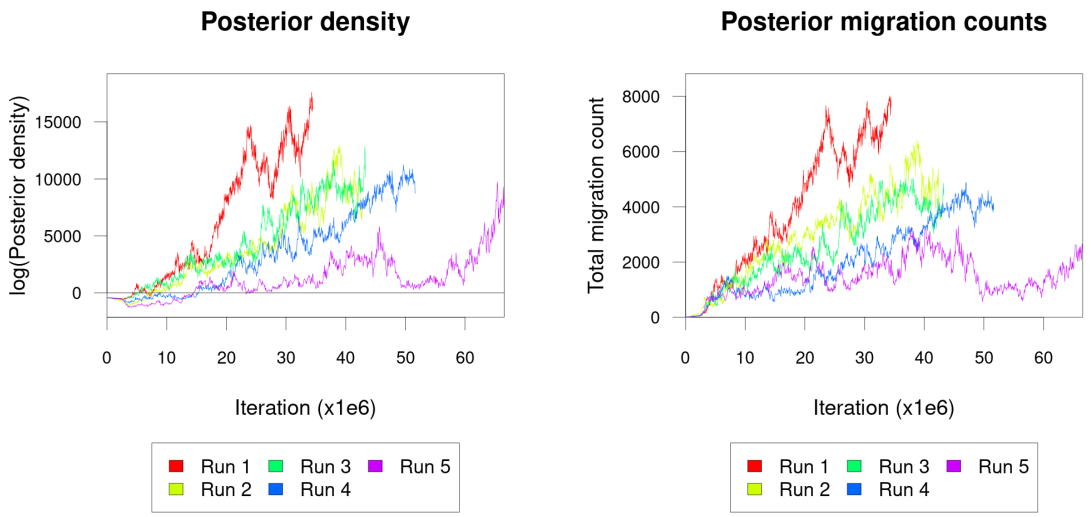


(a) Posterior density evaluations.

(b) Posterior migration counts.

Figure S2: Trace plots of posterior density evaluations and migration counts for the five MultiTypeTree MCMC runs with gamma-distributed priors matching the default Local DTA gamma priors.

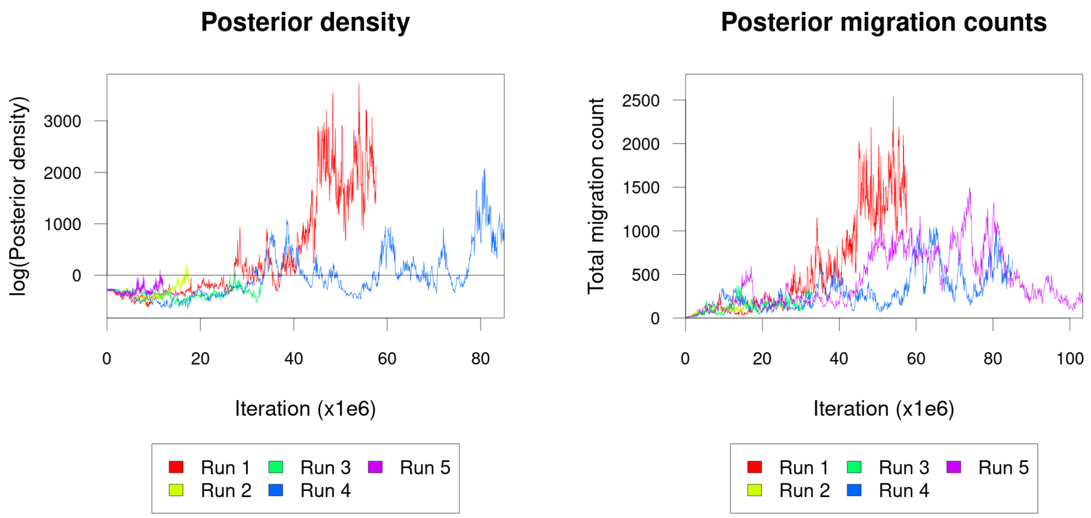


(a) Posterior density evaluations.

(b) Posterior migration counts.

Figure S3: Trace plots of posterior density evaluations and migration counts for the five MultiTypeTree MCMC runs using the default MultiTypeTree lognormal priors.

| | Default priors | | | | | Exp(1) priors | | | | |
|---|---|---|---|---|---|---|---|---|---|---|
| | Run 1 | Run 2 | Run 3 | Run 4 | Run 5 | Run 1 | Run 2 | Run 3 | Run 4 | Run 5 |
| **Coalescent rates** | | | | | | | | | | |
| $\theta_{\text{ANS}}$ | 928 | 1122 | 790 | 1660 | 1001 | 252 | 297 | 203 | 231 | 192 |
| $\theta_{\text{CHA}}$ | 971 | 1510 | 1412 | 1101 | 1673 | 253 | 685 | 713 | 586 | 469 |
| $\theta_{\text{GAL}}$ | 980 | 1279 | 1334 | 1251 | 1280 | 195 | 267 | 162 | 132 | 110 |
| $\theta_{\text{PAS}}$ | 1085 | 1702 | 1083 | 1460 | 1541 | 268 | 90 | 138 | 218 | 252 |
| $\theta_{\text{MEX}}$ | 1482 | 1418 | 1499 | 1039 | 1186 | 43 | 137 | 41 | 60 | 130 |
| **Migration rates** | | | | | | | | | | |
| $\lambda_{\text{CHA,ANS}}$ | 720 | 876 | 1002 | 910 | 866 | 119 | 368 | 356 | 232 | 169 |
| $\lambda_{\text{GAL,ANS}}$ | 983 | 1335 | 1045 | 1355 | 1079 | 78 | 287 | 265 | 260 | 187 |
| $\lambda_{\text{PAS,ANS}}$ | 832 | 735 | 1032 | 728 | 800 | 129 | 156 | 253 | 218 | 99 |
| $\lambda_{\text{MEX,ANS}}$ | 802 | 886 | 1092 | 804 | 868 | 146 | 99 | 176 | 126 | 87 |
| $\lambda_{\text{ANS,CHA}}$ | 287 | 1185 | 1059 | 606 | 1166 | 34 | 109 | 111 | 72 | 199 |
| $\lambda_{\text{GAL,CHA}}$ | 1733 | 1371 | 1413 | 1595 | 1541 | 72 | 104 | 87 | 70 | 57 |
| $\lambda_{\text{PAS,CHA}}$ | 1545 | 1454 | 1695 | 1345 | 1541 | 94 | 49 | 184 | 131 | 79 |
| $\lambda_{\text{MEX,CHA}}$ | 1234 | 1106 | 1687 | 1198 | 1383 | 85 | 57 | 97 | 86 | 86 |
| $\lambda_{\text{ANS,GAL}}$ | 847 | 826 | 976 | 835 | 866 | 109 | 97 | 44 | 48 | 50 |
| $\lambda_{\text{CHA,GAL}}$ | 1008 | 988 | 1012 | 802 | 927 | 82 | 199 | 149 | 107 | 110 |
| $\lambda_{\text{PAS,GAL}}$ | 1196 | 1335 | 1269 | 1555 | 1292 | 98 | 41 | 152 | 78 | 70 |
| $\lambda_{\text{MEX,GAL}}$ | 1620 | 1662 | 971 | 1128 | 1399 | 77 | 54 | 37 | 54 | 90 |
| $\lambda_{\text{ANS,PAS}}$ | 625 | 668 | 838 | 666 | 266 | 99 | 67 | 28 | 55 | 125 |
| $\lambda_{\text{CHA,PAS}}$ | 901 | 1121 | 1137 | 673 | 1155 | 65 | 72 | 82 | 145 | 108 |
| $\lambda_{\text{GAL,PAS}}$ | 1216 | 1459 | 1677 | 1766 | 1262 | 121 | 146 | 127 | 95 | 54 |
| $\lambda_{\text{MEX,PAS}}$ | 1602 | 1262 | 1053 | 1400 | 1632 | 76 | 94 | 105 | 46 | 70 |
| $\lambda_{\text{ANS,MEX}}$ | 648 | 655 | 565 | 507 | 631 | 162 | 82 | 41 | 36 | 71 |
| $\lambda_{\text{CHA,MEX}}$ | 847 | 666 | 717 | 596 | 834 | 74 | 248 | 190 | 125 | 205 |
| $\lambda_{\text{GAL,MEX}}$ | 1149 | 1494 | 1061 | 1240 | 1118 | 74 | 141 | 81 | 52 | 111 |
| $\lambda_{\text{PAS,MEX}}$ | 1447 | 1459 | 1363 | 1204 | 1134 | 107 | 61 | 134 | 90 | 115 |

Table S6: Effective sample size (ESS) results for the AIV analysis with both default gamma-distributed priors and Exp(1) priors.

| | $\theta_x$ | $\lambda_{x,\text{ANS}}$ | $\lambda_{x,\text{CHA}}$ | $\lambda_{x,\text{GAL}}$ | $\lambda_{x,\text{PAS}}$ | $\lambda_{x,\text{MEX}}$ |
|---|---|---|---|---|---|---|
| $x = \text{ANS}$ | 1.0016 | — | **1.0209** | 1.0034 | 1.0120 | 1.0083 |
| $x = \text{CHA}$ | 1.0014 | 1.0005 | — | 1.0016 | 1.0056 | 1.0060 |
| $x = \text{GAL}$ | 1.0021 | 1.0013 | 1.0010 | — | 1.0022 | 1.0010 |
| $x = \text{PAS}$ | 1.0015 | 1.0023 | 1.0017 | 1.0035 | — | 1.0017 |
| $x = \text{MEX}$ | **1.0031** | 1.0014 | 1.0021 | 1.0005 | 1.0013 | — |

(a) $\hat{R}$ results for the default prior AIV analysis.

| | $\theta_x$ | $\lambda_{x,\text{ANS}}$ | $\lambda_{x,\text{CHA}}$ | $\lambda_{x,\text{GAL}}$ | $\lambda_{x,\text{PAS}}$ | $\lambda_{x,\text{MEX}}$ |
|---|---|---|---|---|---|---|
| $x = \text{ANS}$ | 1.2681 | — | 2.2809 | 1.2762 | 1.8498 | **3.6655** |
| $x = \text{CHA}$ | 1.1470 | 1.0708 | — | 1.0141 | 1.8452 | 1.3401 |
| $x = \text{GAL}$ | 1.0772 | 1.2158 | 1.1500 | — | 1.1119 | 1.4620 |
| $x = \text{PAS}$ | **2.2424** | 1.1923 | 1.2051 | 1.0745 | — | 1.2278 |
| $x = \text{MEX}$ | 1.8165 | 1.4964 | 1.2981 | 1.5730 | 1.8347 | — |

(b) $\hat{R}$ results for the Exp(1) AIV analysis.

Table S7: Gelman–Rubin $\hat{R}$ statistics for each evolutionary parater for both AIV analyses. The greatest $\hat{R}$ value is highlighted in **bold**.

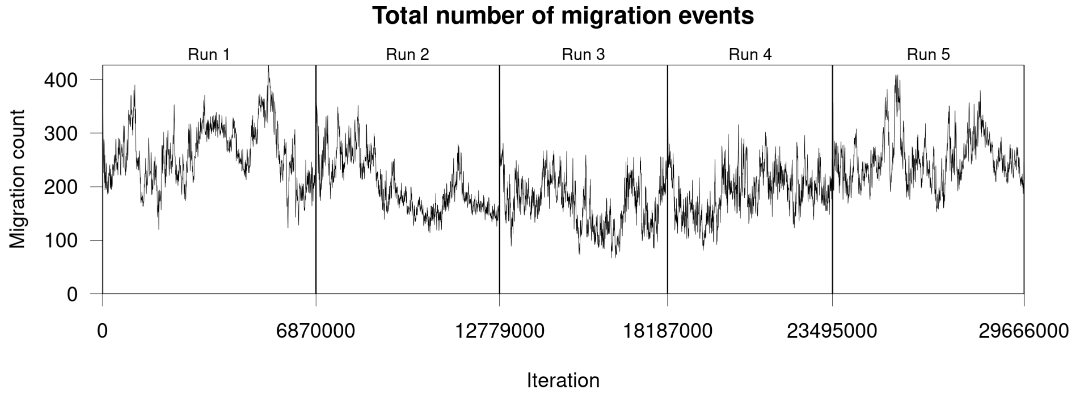


(a) Trace plot of the total migration count for the AIV analysis with Exp(1) priors.

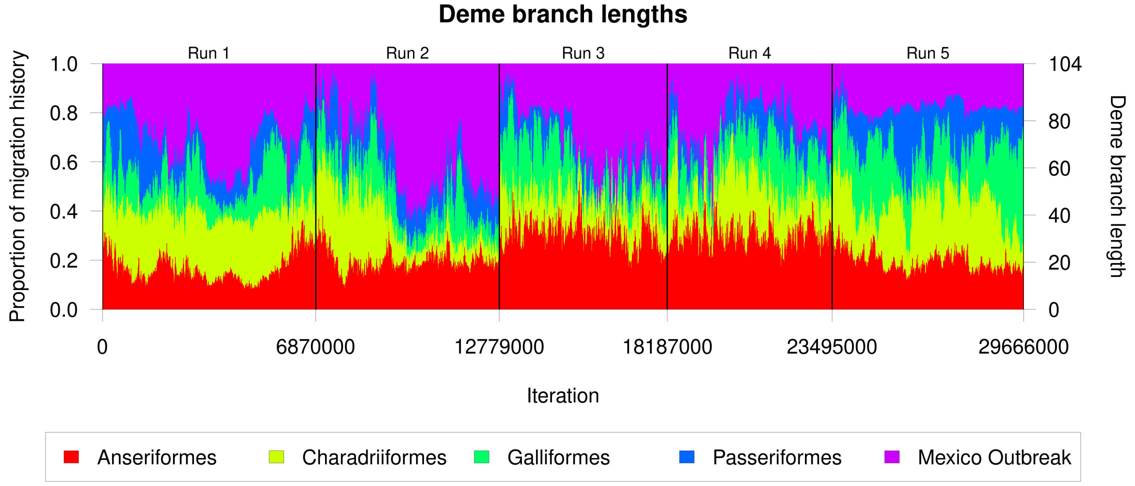


(b) Deme branch lengths for the AIV analysis with Exp(1) priors.

Figure S4: Trace plots of migration history summary statistics for five MCMC runs using Local DTA with Exp(1) priors on an AIV dataset.

| Run ID | $N$ |
|---|---|
| Run 1 | 3687700 |
| Run 2 | 4650900 |
| Run 3 | 4755200 |
| Run 4 | 4411300 |
| Run 5 | 4472400 |
| Run 6 | 4469600 |
| Run 7 | 3021400 |
| Run 8 | 3159500 |
| Run 9 | 3130800 |
| Run 10 | 3924100 |
| Run 11 | 4169800 |

Table S8: Number of iterations in each run for the Cholera analysis.

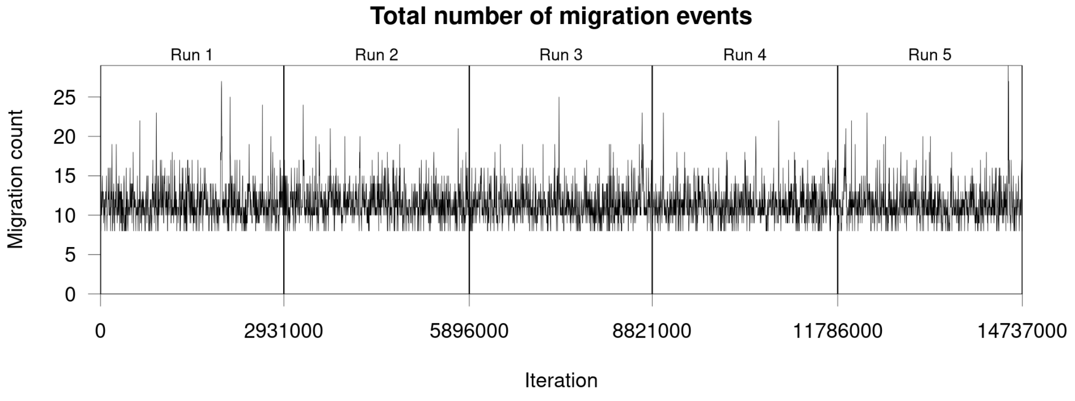


(a) Trace plot of the total migration count for the AIV analysis with default Local DTA priors.

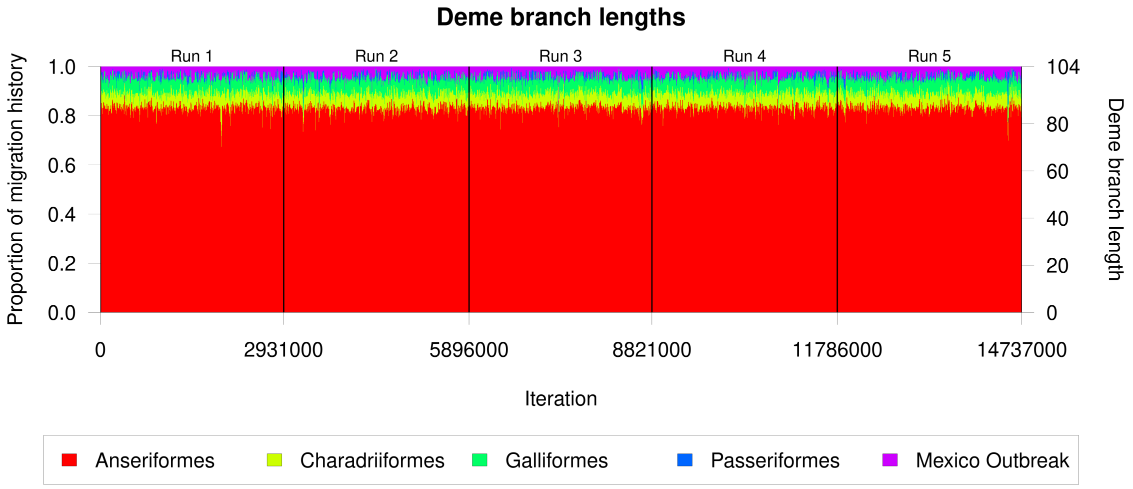


(b) Deme branch lengths for the AIV analysis with default Local DTA priors.

Figure S5: Trace plots of migration history summary statistics for five MCMC runs using default Local DTA priors on an AIV dataset.

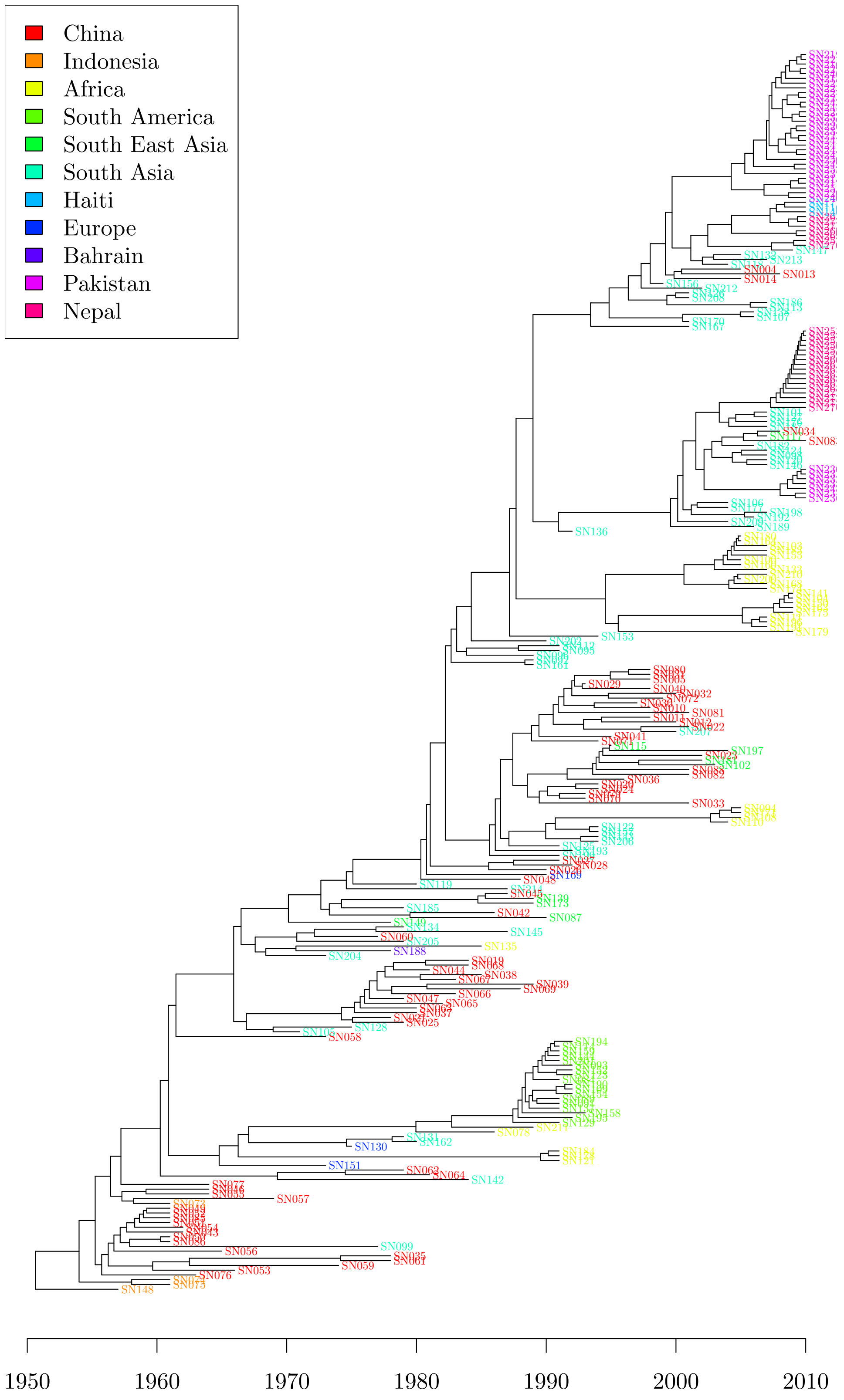


Figure S6: Dated phylogeny used as input of the cholera analysis.

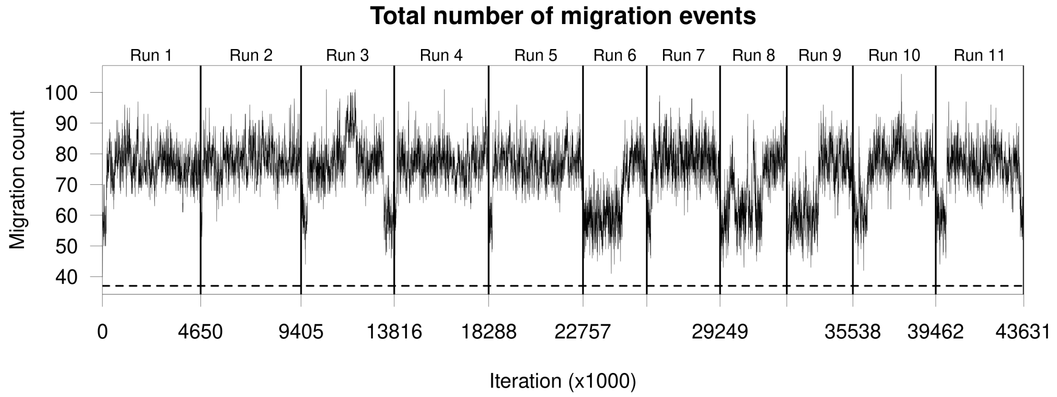


(a) Trace plot of total migration counts. The dashed line shows the number of migration events required in a maximum parsimony migration history (37).

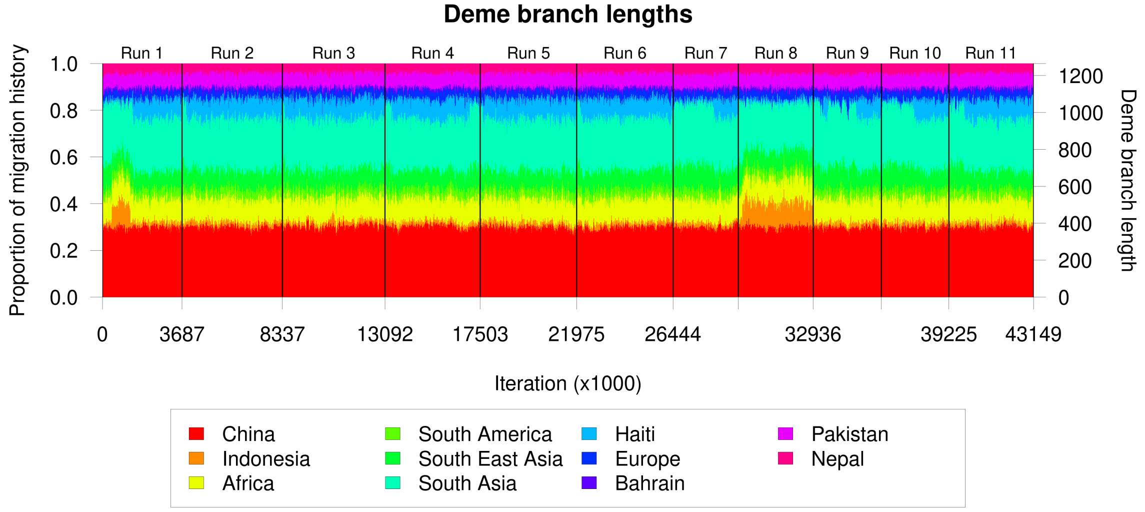


(b) Stacked trace plots of total deme branch lengths.

Figure S7: Summary statistics for mixing over migration histories for the eleven MCMC samples in the preliminary Cholera analysis using default Gamma-distributed priors.

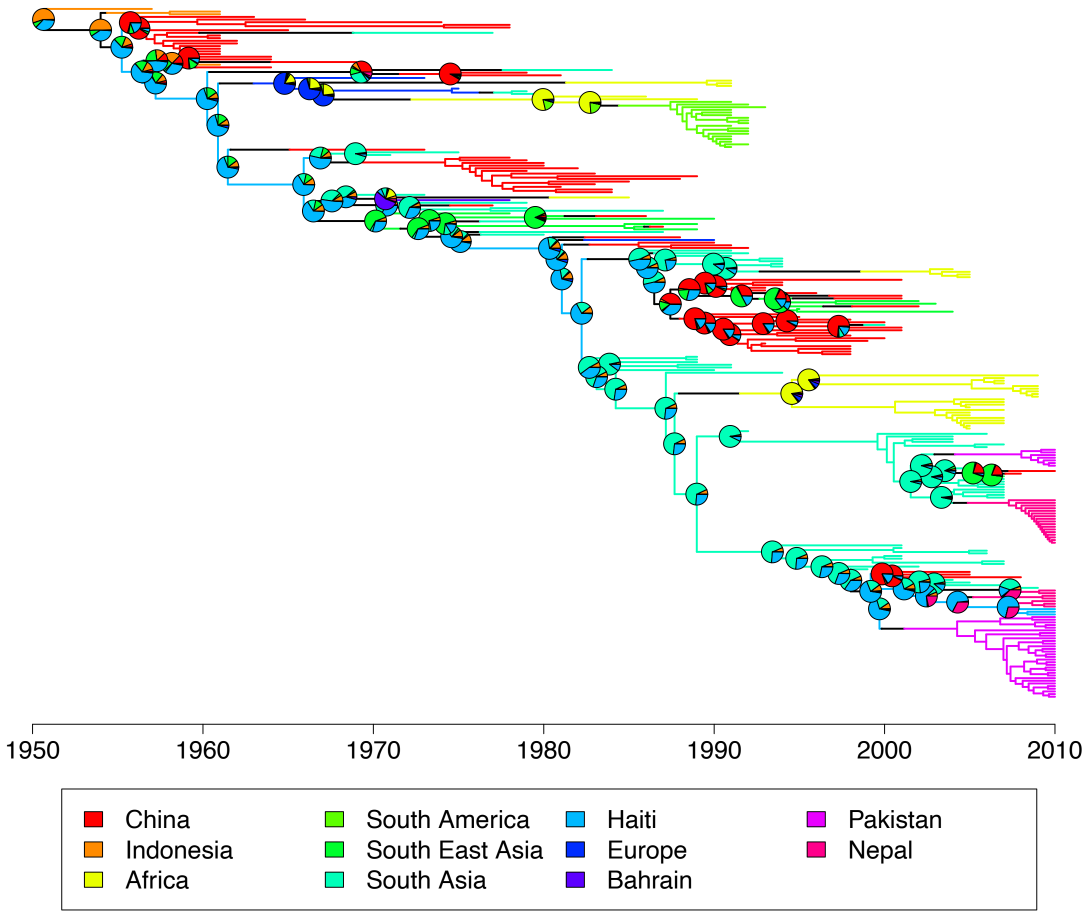


Figure S8: 60% consensus migration history for the eleven MCMC samples in the preliminary Cholera analysis using default Gamma-distributed priors.

| | Run 1 | Run 2 | Run 3 | Run 4 | Run 5 | Run 6 | Run 7 | Run 8 | Run 9 | Run 10 | Run 11 |
|---|---|---|---|---|---|---|---|---|---|---|---|
| **Coalescent rates** | | | | | | | | | | | |
| $\theta_{\mathrm{CHN}}$ | 863 | 1131 | 1275 | 1381 | 1402 | 1074 | **715** | 737 | 842 | 1125 | 1021 |
| $\theta_{\mathrm{IDN}}$ | **307** | 1376 | 1201 | 1002 | 1145 | 1178 | 948 | 883 | 996 | 1218 | 1291 |
| $\theta_{\mathrm{AFR}}$ | 601 | 781 | 674 | 609 | 656 | 763 | 659 | **520** | 716 | 574 | 801 |
| $\theta_{\mathrm{SA}}$ | 1931 | 1947 | 2630 | 1895 | 2024 | 1959 | 1378 | 1504 | **1001** | 2133 | 1707 |
| $\theta_{\mathrm{IC}}$ | 587 | 811 | 894 | 564 | 814 | 884 | 419 | **339** | 459 | 569 | 487 |
| $\theta_{\mathrm{SAS}}$ | 745 | 705 | 1050 | 478 | 789 | 1000 | 519 | **436** | 453 | 706 | 895 |
| $\theta_{\mathrm{HTI}}$ | 98 | 182 | 205 | 148 | 213 | 182 | **88** | 92 | 112 | 124 | 139 |
| $\theta_{\mathrm{EUR}}$ | 971 | 1067 | 993 | 718 | 921 | 1274 | 578 | **301** | 433 | 414 | 634 |
| $\theta_{\mathrm{BHR}}$ | 1150 | 1500 | 1299 | 1678 | 1878 | 1784 | 1487 | 1052 | 1351 | **876** | 1347 |
| $\theta_{\mathrm{PAK}}$ | 1915 | 2470 | 2163 | 2035 | 2146 | 2404 | **1373** | 1545 | 1663 | 2458 | 2324 |
| $\theta_{\mathrm{NPL}}$ | 1424 | 2130 | 2145 | 1781 | 2070 | 1789 | 1249 | 1250 | **1032** | 1516 | 1428 |
| **Coalescent rates** | | | | | | | | | | | |
| $\lambda_{\mathrm{IDN,CHN}}$ | 1110 | 1287 | 1149 | 1661 | 1366 | 1247 | 971 | 990 | **921** | 1013 | 1186 |
| $\lambda_{\mathrm{AFR,CHN}}$ | 1624 | 1505 | 1811 | 2293 | 1279 | 1723 | 1092 | 1383 | **1014** | 1533 | 1791 |
| $\lambda_{\mathrm{SA,CHN}}$ | 1554 | 2039 | 1882 | 2211 | 1572 | 1461 | 1337 | **1249** | 1454 | 1306 | 1594 |
| $\lambda_{\mathrm{IC,CHN}}$ | 819 | 841 | 799 | 851 | 819 | 767 | 534 | **469** | 611 | 569 | 651 |
| $\lambda_{\mathrm{SAS,CHN}}$ | 835 | 754 | 978 | 633 | 818 | 649 | 684 | 534 | 565 | 601 | **518** |
| $\lambda_{\mathrm{HTI,CHN}}$ | 1160 | 1975 | 1582 | 1644 | 1670 | 1249 | 961 | 1142 | **872** | 1129 | 1202 |
| $\lambda_{\mathrm{EUR,CHN}}$ | 1357 | 1611 | 1155 | 1426 | 2105 | 1553 | **826** | 1393 | 942 | 1440 | 1399 |
| $\lambda_{\mathrm{BHR,CHN}}$ | 1790 | 2273 | 2365 | 1801 | 1672 | 1719 | 1538 | 1653 | 1521 | 1705 | **1477** |
| $\lambda_{\mathrm{PAK,CHN}}$ | 1408 | 1874 | 1740 | 1717 | 1709 | 1752 | 1076 | 1286 | **902** | 1996 | 1641 |
| $\lambda_{\mathrm{NPL,CHN}}$ | 1297 | 1719 | 1709 | 1580 | 1617 | 1116 | **928** | 1148 | 1293 | 1551 | 1457 |
| $\lambda_{\mathrm{CHN,IDN}}$ | **39** | 701 | 621 | 819 | 270 | 456 | 383 | 316 | 343 | 387 | 693 |
| $\lambda_{\mathrm{AFR,IDN}}$ | 634 | 587 | **360** | 636 | 970 | 741 | 483 | 570 | 625 | 587 | 1030 |
| $\lambda_{\mathrm{SA,IDN}}$ | 1350 | 3259 | 1819 | 2042 | 2005 | 1877 | 1303 | 1869 | **1178** | 2278 | 1336 |
| $\lambda_{\mathrm{IC,IDN}}$ | **163** | 1206 | 1077 | 1585 | 997 | 1162 | 750 | 966 | 1162 | 718 | 775 |
| $\lambda_{\mathrm{SAS,IDN}}$ | **27** | 1262 | 1266 | 922 | 978 | 724 | 677 | 590 | 781 | 937 | 933 |
| $\lambda_{\mathrm{HTI,IDN}}$ | **752** | 1211 | 1231 | 918 | 834 | 951 | 944 | 1290 | 956 | 1020 | 1129 |
| $\lambda_{\mathrm{EUR,IDN}}$ | **392** | 1327 | 1412 | 1238 | 1443 | 1299 | 695 | 873 | 1019 | 1584 | 1629 |
| $\lambda_{\mathrm{BHR,IDN}}$ | **938** | 1959 | 1838 | 1587 | 2223 | 1581 | 953 | 1719 | 1173 | 1814 | 1690 |
| $\lambda_{\mathrm{PAK,IDN}}$ | **671** | 1862 | 1770 | 1330 | 1763 | 1654 | 1106 | 1817 | 1325 | 2348 | 1698 |
| $\lambda_{\mathrm{NPL,IDN}}$ | **516** | 1835 | 2052 | 2030 | 1938 | 1423 | 1017 | 1185 | 1102 | 1314 | 2181 |
| $\lambda_{\mathrm{CHN,AFR}}$ | 1539 | 1684 | 1615 | 1726 | 1698 | 1679 | 1235 | 1452 | **1102** | 1477 | 1441 |

| | | | | | | | | | | | |
|---|---|---|---|---|---|---|---|---|---|---|---|
| $\lambda_{\text{IDN,AFR}}$ | 1486 | 2006 | 1928 | 1703 | 1765 | 1922 | **1015** | 1147 | 1378 | 2004 | 1383 |
| $\lambda_{\text{SA,AFR}}$ | 1200 | 1797 | 1413 | 1588 | 1271 | 1479 | 1157 | 1190 | **909** | 1226 | 1187 |
| $\lambda_{\text{IC,AFR}}$ | 1670 | 1864 | 1847 | 1678 | 1951 | 2446 | 1331 | **1254** | 1680 | 1778 | 1292 |
| $\lambda_{\text{SAS,AFR}}$ | 1672 | 1709 | 2599 | 1807 | 1890 | 1562 | **985** | 1626 | 1481 | 1621 | 1731 |
| $\lambda_{\text{HTI,AFR}}$ | 1501 | 1982 | 2233 | 1698 | 2309 | 1999 | **834** | 1456 | 1025 | 1447 | 1771 |
| $\lambda_{\text{EUR,AFR}}$ | 637 | 895 | 839 | 777 | 917 | 598 | **510** | 608 | 577 | 660 | 875 |
| $\lambda_{\text{BHR,AFR}}$ | 1233 | 1607 | 1363 | 1490 | 1264 | 1318 | **1071** | 1089 | **1071** | 1348 | 1880 |
| $\lambda_{\text{PAK,AFR}}$ | 1859 | 1630 | 2547 | 1824 | 2008 | 1817 | 1628 | **1289** | 1362 | 1578 | 1742 |
| $\lambda_{\text{NPL,AFR}}$ | 1794 | 1910 | 2183 | 1905 | 2046 | 2060 | **1165** | 1527 | 1657 | 1427 | 1884 |
| $\lambda_{\text{CHN,SA}}$ | 1177 | 2349 | 2236 | 1755 | 1538 | 1814 | 1371 | 1322 | 1117 | 1084 | **805** |
| $\lambda_{\text{IDN,SA}}$ | **1139** | 2052 | 1845 | 2183 | 1838 | 1776 | 1411 | 1595 | 1252 | 1606 | 1634 |
| $\lambda_{\text{AFR,SA}}$ | 854 | 661 | 731 | 709 | 631 | 505 | **450** | 534 | 476 | 495 | 626 |
| $\lambda_{\text{IC,SA}}$ | 1853 | 2046 | 2194 | 1897 | 1883 | 2525 | 1712 | 1250 | 1246 | 1331 | **1141** |
| $\lambda_{\text{SAS,SA}}$ | 998 | 2042 | 1840 | 1967 | 1616 | 1153 | **782** | 1121 | 1038 | 1286 | 1781 |
| $\lambda_{\text{HTI,SA}}$ | 1784 | 1845 | 2107 | 2195 | 2036 | 1781 | 1312 | **1202** | 1413 | 1745 | 1563 |
| $\lambda_{\text{EUR,SA}}$ | 1205 | 1546 | 1021 | 1201 | 1428 | 1015 | **917** | 1141 | 1135 | 949 | 1601 |
| $\lambda_{\text{BHR,SA}}$ | 1792 | 2066 | 1765 | 2275 | 1931 | 1711 | 1597 | 1349 | **1257** | 2116 | 1671 |
| $\lambda_{\text{PAK,SA}}$ | 1577 | 1836 | 2354 | 1640 | 1969 | 2148 | 1807 | 1435 | **1394** | 2328 | 2227 |
| $\lambda_{\text{NPL,SA}}$ | 1862 | 2112 | 1743 | 1662 | 1446 | 1740 | **1221** | 1492 | 1593 | 1500 | 1772 |
| $\lambda_{\text{CHN,IC}}$ | 159 | 246 | 267 | 211 | 238 | 258 | **136** | 147 | 177 | 182 | 180 |
| $\lambda_{\text{IDN,IC}}$ | 480 | 629 | 781 | 680 | 932 | 900 | 363 | 316 | **296** | 589 | 547 |
| $\lambda_{\text{AFR,IC}}$ | 1541 | 2091 | 1608 | 1775 | 1430 | 1492 | 1289 | **1175** | 1601 | 1445 | 1779 |
| $\lambda_{\text{SA,IC}}$ | 1550 | 2290 | 2090 | 2062 | 1331 | 1479 | **942** | 1166 | 1371 | 2219 | 2007 |
| $\lambda_{\text{SAS,IC}}$ | 311 | 703 | 663 | 324 | 700 | 430 | 194 | **191** | 224 | 278 | 250 |
| $\lambda_{\text{HTI,IC}}$ | 763 | 783 | 630 | 781 | 657 | 923 | 740 | 709 | 618 | 756 | **558** |
| $\lambda_{\text{EUR,IC}}$ | 1169 | 1690 | 1882 | 833 | 1347 | 1336 | **525** | 668 | 946 | 756 | 843 |
| $\lambda_{\text{BHR,IC}}$ | 1565 | 2349 | 1662 | 1879 | 1698 | 1566 | 948 | 1436 | **746** | 1581 | 1423 |
| $\lambda_{\text{PAK,IC}}$ | 1294 | 2305 | 1766 | 2339 | 1807 | 2411 | **1162** | 1368 | 1347 | 1651 | 1897 |
| $\lambda_{\text{NPL,IC}}$ | 1195 | 1632 | 2148 | 1513 | 1266 | 1575 | **1100** | 1707 | 1251 | 1457 | 1473 |
| $\lambda_{\text{CHN,SAS}}$ | 216 | 267 | 344 | 221 | 294 | 260 | **150** | 181 | 155 | 225 | 221 |
| $\lambda_{\text{IDN,SAS}}$ | **617** | 1074 | 1072 | 1169 | 1397 | 1676 | 890 | 1311 | 1095 | 994 | 1571 |
| $\lambda_{\text{AFR,SAS}}$ | 962 | 972 | 1140 | 820 | 851 | 973 | **614** | 706 | 722 | 675 | 1121 |
| $\lambda_{\text{SA,SAS}}$ | 1601 | 1775 | 2005 | 1783 | 1709 | 1755 | **1217** | 1275 | 1427 | 1698 | 1837 |
| $\lambda_{\text{IC,SAS}}$ | 575 | 597 | 806 | 633 | 762 | 835 | **367** | 516 | 476 | 486 | 595 |
| $\lambda_{\text{HTI,SAS}}$ | 754 | 601 | 540 | 598 | 654 | 621 | 692 | 667 | **520** | 541 | 955 |
| $\lambda_{\text{EUR,SAS}}$ | 877 | 1055 | 1172 | 686 | 1035 | 800 | 534 | **476** | 553 | 644 | 895 |
| $\lambda_{\text{BHR,SAS}}$ | 991 | 1064 | 1378 | 1014 | 1333 | 1146 | 837 | 856 | **828** | 1044 | 839 |

| | | | | | | | | | | | |
|---|---|---|---|---|---|---|---|---|---|---|---|
| $\lambda_{PAK,SAS}$ | 913 | 1210 | 1704 | 837 | 1254 | 1636 | **568** | 596 | 670 | 939 | 735 |
| $\lambda_{NPL,SAS}$ | **604** | 1088 | 1214 | 969 | 1102 | 1157 | 805 | 721 | 800 | 757 | 1179 |
| $\lambda_{CHN,HTI}$ | 51 | 126 | 176 | 90 | 131 | 118 | **34** | 43 | 43 | 83 | 83 |
| $\lambda_{IDN,HTI}$ | 207 | 445 | 462 | 341 | 417 | 353 | 211 | 195 | **179** | 297 | 306 |
| $\lambda_{AFR,HTI}$ | 677 | 826 | 735 | 822 | 739 | 644 | **479** | 707 | 494 | 761 | 947 |
| $\lambda_{SA,HTI}$ | 1133 | 1517 | 1315 | 1126 | 1272 | 1336 | 902 | 1457 | 1223 | 995 | **810** |
| $\lambda_{IC,HTI}$ | 168 | 393 | 428 | 259 | 413 | 372 | **157** | 167 | 170 | 221 | 254 |
| $\lambda_{SAS,HTI}$ | 52 | 135 | 216 | 56 | 149 | 144 | 31 | **28** | 30 | 62 | 104 |
| $\lambda_{EUR,HTI}$ | 344 | 918 | 706 | 437 | 718 | 670 | 299 | **287** | 331 | 415 | 522 |
| $\lambda_{BHR,HTI}$ | 624 | 804 | 777 | 1006 | 904 | 794 | **363** | 415 | 689 | 487 | 826 |
| $\lambda_{PAK,HTI}$ | 710 | 1096 | 1039 | 751 | 968 | 929 | 629 | 508 | **385** | 725 | 684 |
| $\lambda_{NPL,HTI}$ | 246 | 599 | 844 | 351 | 533 | 586 | 227 | **206** | 235 | 358 | 432 |
| $\lambda_{CHN,EUR}$ | 1177 | 1539 | 1208 | 375 | 1185 | 1367 | 202 | **55** | 121 | 91 | 946 |
| $\lambda_{IDN,EUR}$ | 1324 | 1610 | 1892 | 1268 | 1868 | 1472 | 913 | **530** | 897 | 712 | 1435 |
| $\lambda_{AFR,EUR}$ | 502 | 587 | 519 | **290** | 438 | 575 | 466 | 391 | 397 | 314 | 509 |
| $\lambda_{SA,EUR}$ | 1127 | 1674 | 1324 | 1361 | 1173 | 1484 | 1395 | **929** | 995 | 1239 | 1309 |
| $\lambda_{IC,EUR}$ | 1360 | 2058 | 2463 | 1863 | 1941 | 2164 | 899 | **573** | 615 | 1146 | 1527 |
| $\lambda_{SAS,EUR}$ | 984 | 1789 | 1331 | 755 | 1946 | 1360 | 249 | **62** | 314 | 174 | 823 |
| $\lambda_{HTI,EUR}$ | 1228 | 2225 | 2051 | 1354 | 1789 | 1672 | 1465 | 981 | **871** | 1184 | 1538 |
| $\lambda_{BHR,EUR}$ | 1538 | 1710 | 1619 | 2009 | 2204 | 1863 | 1379 | 959 | **934** | 1671 | 2056 |
| $\lambda_{PAK,EUR}$ | 1773 | 1788 | 1883 | 1499 | 1606 | 2065 | 1140 | **978** | 1425 | 1419 | 1280 |
| $\lambda_{NPL,EUR}$ | 1334 | 1568 | 2312 | 2070 | 1609 | 1530 | 1009 | **315** | 811 | 558 | 1130 |
| $\lambda_{CHN,BHR}$ | 1006 | 1016 | 1333 | 1200 | 1333 | 1205 | **593** | 848 | 947 | 915 | 1230 |
| $\lambda_{IDN,BHR}$ | 1528 | 1816 | 1825 | 2453 | 2027 | 1844 | **1257** | 1612 | 1736 | 2040 | 1695 |
| $\lambda_{AFR,BHR}$ | **395** | 730 | 613 | 826 | 602 | 548 | 592 | 431 | 495 | 430 | 447 |
| $\lambda_{SA,BHR}$ | 1719 | 2409 | 2030 | 2461 | 1764 | 1739 | 1392 | 1398 | **1262** | 2274 | 1710 |
| $\lambda_{IC,BHR}$ | **1224** | 1851 | 2092 | 1504 | 1509 | 1374 | 1335 | 1273 | 1321 | 1536 | 1669 |
| $\lambda_{SAS,BHR}$ | 809 | 1277 | 1057 | 1645 | 1628 | 1354 | **682** | 1092 | 1097 | 743 | 1399 |
| $\lambda_{HTI,BHR}$ | 1670 | 2116 | 2921 | 2173 | 1728 | 1724 | **931** | 1251 | 1110 | 1486 | 1433 |
| $\lambda_{EUR,BHR}$ | 1426 | 1812 | 1448 | 1178 | 1232 | 1165 | **955** | 1139 | 1139 | 1639 | 1366 |
| $\lambda_{PAK,BHR}$ | 1536 | 1971 | 1559 | 2091 | 1923 | 1863 | **1037** | 1164 | 1185 | 1518 | 1794 |
| $\lambda_{NPL,BHR}$ | **1111** | 1543 | 2582 | 1872 | 2894 | 1848 | 1528 | 1217 | 1181 | 1437 | 1227 |
| $\lambda_{CHN,PAK}$ | 1172 | 1966 | 1260 | 1471 | 1275 | 1189 | 917 | 1047 | **607** | 1100 | 1267 |
| $\lambda_{IDN,PAK}$ | **1058** | 2578 | 2160 | 1964 | 2168 | 1532 | 1148 | 1488 | 1246 | 1935 | 1579 |
| $\lambda_{AFR,PAK}$ | 916 | 1337 | 1180 | 1161 | 813 | 1203 | 1076 | 1013 | **787** | 1510 | 1240 |
| $\lambda_{SA,PAK}$ | 1352 | 2757 | 1999 | 2124 | 1873 | 1607 | 2259 | **1292** | 1718 | 2010 | 2382 |
| $\lambda_{IC,PAK}$ | 1894 | 1784 | 1748 | 1674 | 1772 | 1432 | 1539 | **1082** | 1085 | 1365 | 1589 |

| | | | | | | | | | | | |
|---|---|---|---|---|---|---|---|---|---|---|---|
| $\lambda_{\mathrm{SAS,PAK}}$ | 1236 | 1434 | 1389 | 1126 | 1353 | 1038 | 1026 | 909 | **741** | 1389 | 855 |
| $\lambda_{\mathrm{HTI,PAK}}$ | 1371 | 1722 | 2679 | 1845 | 2148 | 1836 | 1426 | 1335 | **916** | 2189 | 1854 |
| $\lambda_{\mathrm{EUR,PAK}}$ | 1829 | 2071 | 1295 | 1622 | 1738 | 1450 | **953** | 1448 | 1051 | 1493 | 1806 |
| $\lambda_{\mathrm{BHR,PAK}}$ | 1574 | 1668 | 1881 | 1796 | 2137 | 2035 | **1217** | 1359 | 1609 | 1411 | 1821 |
| $\lambda_{\mathrm{NPL,PAK}}$ | 1894 | 1906 | 2236 | 2155 | 1462 | 1989 | 1232 | 1163 | **1092** | 2164 | 1858 |
| $\lambda_{\mathrm{CHN,NPL}}$ | 1172 | 856 | 1066 | 953 | 780 | 835 | 958 | **632** | 738 | 812 | 1072 |
| $\lambda_{\mathrm{IDN,NPL}}$ | 1345 | 1794 | 1921 | 2015 | 2079 | 1715 | **1170** | 1583 | 1449 | 1399 | 1586 |
| $\lambda_{\mathrm{AFR,NPL}}$ | 738 | 760 | **573** | 1036 | 786 | 1041 | 838 | 632 | 1074 | 1023 | 1128 |
| $\lambda_{\mathrm{SA,NPL}}$ | 1928 | 2379 | 2119 | 1620 | 1680 | 1858 | 1541 | 1266 | **975** | 1835 | 1576 |
| $\lambda_{\mathrm{IC,NPL}}$ | 1134 | 1126 | 1436 | 1162 | 1474 | 964 | 1045 | 1453 | **837** | 1157 | 1416 |
| $\lambda_{\mathrm{SAS,NPL}}$ | 567 | 687 | 686 | 857 | 551 | 612 | 634 | 526 | **459** | 547 | 718 |
| $\lambda_{\mathrm{HTI,NPL}}$ | 533 | 939 | 1268 | 766 | 1018 | 994 | **465** | 495 | 522 | 601 | 720 |
| $\lambda_{\mathrm{EUR,NPL}}$ | 1496 | 1488 | 1440 | 1178 | 967 | 1058 | 1042 | **811** | 899 | 1335 | 1092 |
| $\lambda_{\mathrm{BHR,NPL}}$ | 1388 | 1808 | 2163 | 2190 | 1846 | 2288 | 1586 | **1033** | 1091 | 1485 | 1792 |
| $\lambda_{\mathrm{PAK,NPL}}$ | 1334 | 1884 | 2427 | 1954 | 1759 | 1533 | 1322 | 1572 | **1060** | 2069 | 1753 |

Table S9: Effective sample size estimates for each evolutionary parameter in the cholera analysis.

| | $\lambda_{\mathrm{CHN},x}$ | $\lambda_{\mathrm{IDN},x}$ | $\lambda_{\mathrm{AFR},x}$ | $\lambda_{\mathrm{SA},x}$ | $\lambda_{\mathrm{IC},x}$ | $\lambda_{\mathrm{SAS},x}$ |
|---|---|---|---|---|---|---|
| $x$=CHN | — | 1.0026 (1.0046) | 1.0006 (1.0008) | 1.0031 (1.0053) | 1.0146 (1.0286) | 1.0262 (1.0508) |
| $x$=IDN | 1.0008 (1.0013) | — | 1.0009 (1.0012) | 1.0007 (1.0010) | 1.0023 (1.0038) | 1.0030 (1.0053) |
| $x$=AFR | 1.0014 (1.0025) | 1.0045 (1.0072) | — | 1.0036 (1.0064) | 1.0033 (1.0058) | 1.0073 (1.0146) |
| $x$=SA | 1.0010 (1.0016) | 1.0003 (1.0006) | 1.0020 (1.0034) | — | 1.0005 (1.0009) | 1.0011 (1.0019) |
| $x$=IC | 1.0102 (1.0194) | 1.0026 (1.0045) | 1.0016 (1.0025) | 1.0009 (1.0015) | — | 1.0032 (1.0057) |
| $x$=SAS | 1.0043 (1.0087) | 1.0007 (1.0010) | 1.0018 (1.0033) | 1.0015 (1.0024) | 1.0168 (1.0290) | — |
| $x$=HTI | 1.0023 (1.0036) | 1.0016 (1.0031) | 1.0011 (1.0019) | 1.0017 (1.0027) | 1.0043 (1.0085) | 1.0287 (1.0569) |
| $x$=EUR | 1.0012 (1.0023) | 1.0010 (1.0018) | 1.0027 (1.0050) | 1.0019 (1.0030) | 1.0009 (1.0016) | 1.0044 (1.0074) |
| $x$=BHR | 1.0007 (1.0012) | 1.0013 (1.0020) | 1.0014 (1.0024) | 1.0006 (1.0011) | 1.0006 (1.0009) | 1.0024 (1.0044) |
| $x$=PAK | 1.0015 (1.0027) | 1.0018 (1.0030) | 1.0014 (1.0023) | 1.0017 (1.0030) | 1.0010 (1.0015) | 1.0013 (1.0025) |
| $x$=NPL | 1.0021 (1.0037) | 1.0015 (1.0025) | 1.0007 (1.0013) | 1.0007 (1.0011) | 1.0015 (1.0025) | 1.0010 (1.0017) |

| | $\lambda_{\mathrm{HTI},x}$ | $\lambda_{\mathrm{EUR},x}$ | $\lambda_{\mathrm{BHR},x}$ | $\lambda_{\mathrm{PAK},x}$ | $\lambda_{\mathrm{NPL},x}$ | $\theta_x$ |
|---|---|---|---|---|---|---|
| $x$=CHN | **1.0857 (1.1684)** | 1.0053 (1.0069) | 1.0038 (1.0071) | 1.0036 (1.0054) | 1.0024 (1.0042) | 1.0050 (1.0100) |
| $x$=IDN | 1.0055 (1.0110) | 1.0003 (1.0006) | 1.0003 (1.0006) | 1.0011 (1.0019) | 1.0017 (1.0026) | 1.0016 (1.0031) |
| $x$=AFR | 1.0075 (1.0139) | 1.0067 (1.0111) | 1.0023 (1.0039) | 1.0014 (1.0022) | 1.0009 (1.0015) | 1.0024 (1.0047) |
| $x$=SA | 1.0018 (1.0031) | 1.0022 (1.0040) | 1.0004 (1.0006) | 1.0012 (1.0019) | 1.0016 (1.0027) | 1.0009 (1.0018) |
| $x$=IC | 1.0165 (1.0329) | 1.0010 (1.0017) | 1.0007 (1.0013) | 1.0007 (1.0012) | 1.0011 (1.0018) | 1.0026 (1.0050) |
| $x$=SAS | 1.0805 (1.1619) | 1.0024 (1.0036) | 1.0026 (1.0041) | 1.0010 (1.0017) | 1.0038 (1.0067) | 1.0089 (1.0178) |
| $x$=HTI | — | 1.0009 (1.0016) | 1.0013 (1.0024) | 1.0009 (1.0015) | 1.0061 (1.0112) | 1.0219 (1.0424) |
| $x$=EUR | 1.0022 (1.0044) | — | 1.0013 (1.0023) | 1.0013 (1.0018) | 1.0034 (1.0066) | 1.0006 (1.0010) |
| $x$=BHR | 1.0096 (1.0179) | 1.0005 (1.0007) | — | 1.0011 (1.0018) | 1.0022 (1.0033) | 1.0011 (1.0018) |
| $x$=PAK | 1.0027 (1.0051) | 1.0017 (1.0028) | 1.0010 (1.0017) | — | 1.0006 (1.0012) | 1.0010 (1.0019) |
| $x$=NPL | 1.0058 (1.0116) | 1.0010 (1.0017) | 1.0006 (1.0009) | 1.0003 (1.0006) | — | 1.0011 (1.0021) |

Table S10: Gelman–Rubin $\hat{R}$ estimates for each evolutionary parameter in the cholera analysis. Quantities in brakcets give the upper 95% confidence interval for the $\hat{R}$ value.

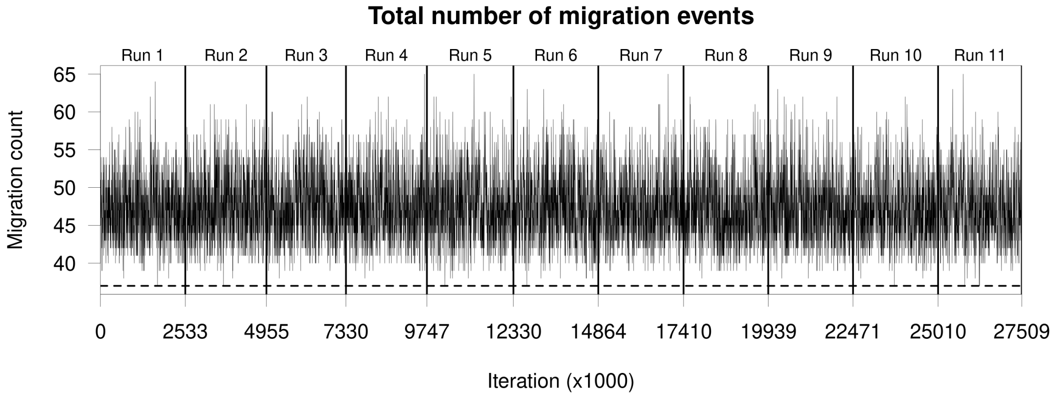


(a) Trace plot of total migration counts. The dashed line shows the number of migration events required in a maximum parsimony migration history (37).

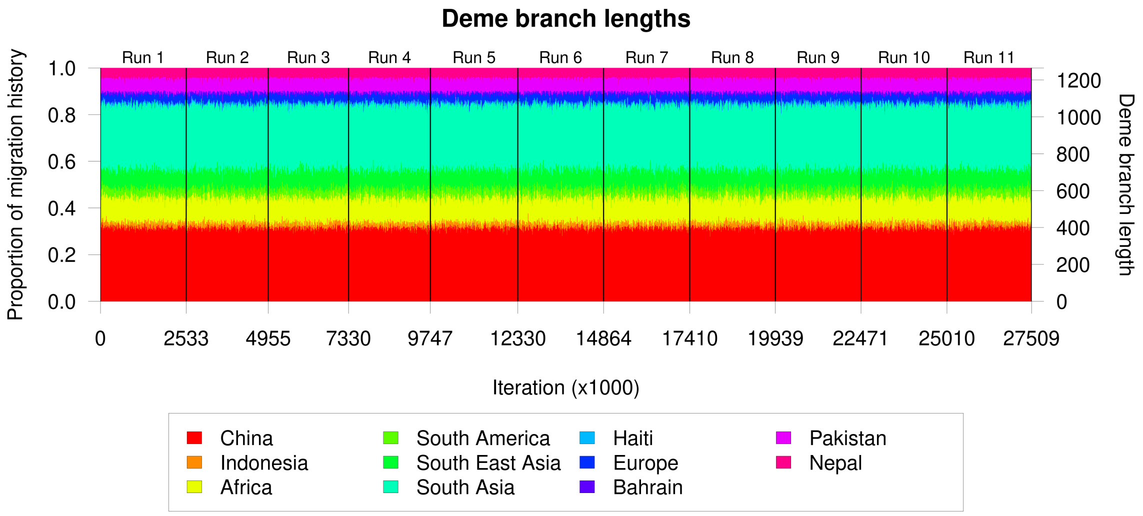


(b) Stacked trace plots of total deme branch lengths.

Figure S9: Summary statistics for mixing over migration histories for the eleven MCMC samples in the Cholera analysis.

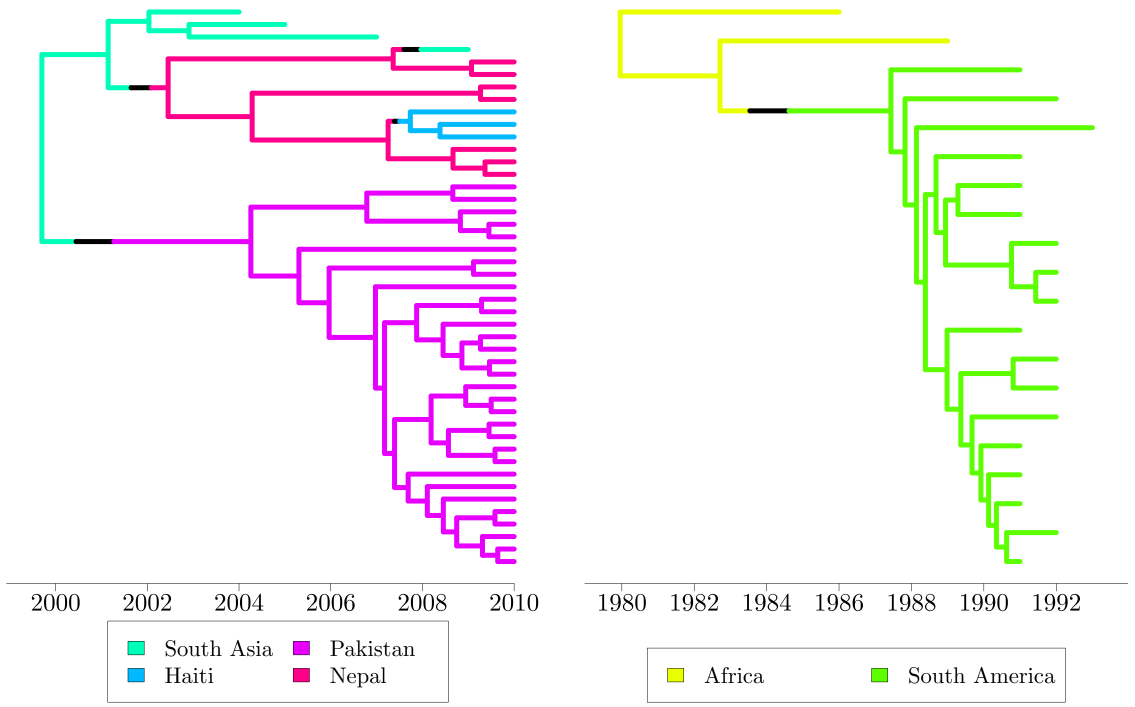


(a) 60% consensus migration history focussing on recent migration events into Haiti.

(b) 60% consensus migration history focussing on migration events into South America.

Figure S10: 60% consensus migration histories.

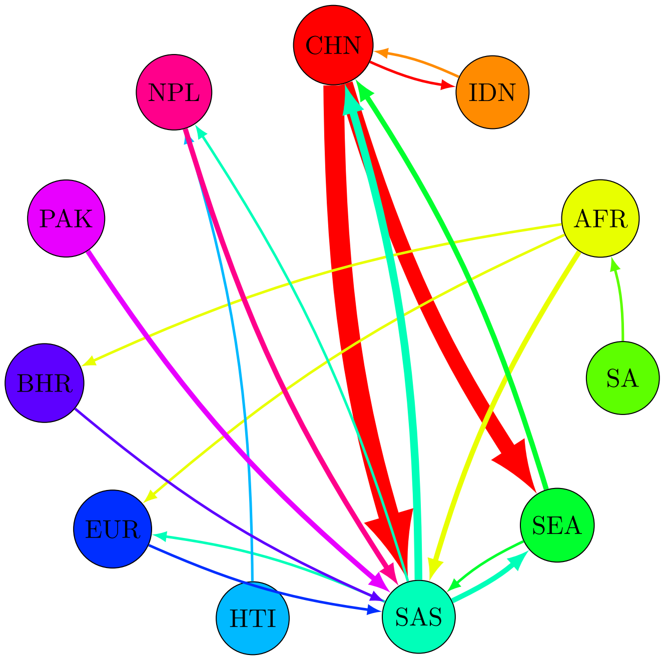


Figure S11: Median number of **backwards-in-time** migration events between pairs of demes in our Cholera analysis. The width of each arrow denotes the relative frequency with which that migration type was observed.
